# Interdigitating Modules for Visual Processing During Locomotion and Rest in Mouse V1

**DOI:** 10.1101/2025.02.21.639505

**Authors:** AM Meier, RD D’Souza, W Ji, EB Han, A Burkhalter

**Affiliations:** Department of Neuroscience, Washington University School of Medicine, St. Louis, MO 63110; USA

**Keywords:** Mouse, visual cortex, layer 1, modular architecture, looped cortical circuits, calcium imaging, visual responses, modulation by locomotion

## Abstract

Layer 1 of V1 has been shown to receive locomotion-related signals from the dorsal lateral geniculate (dLGN) and lateral posterior (LP) thalamic nuclei (***Roth et al., 2016***). Inputs from the dLGN terminate in M2+ patches while inputs from LP target M2− interpatches (***D’Souza et al., 2019***) suggesting that motion related signals are processed in distinct networks. Here, we investigated by calcium imaging in head-fixed awake mice whether L2/3 neurons underneath L1 M2+ and M2− modules are differentially activated by locomotion, and whether distinct networks of feedback connections from higher cortical areas to L1 may contribute to these differences. We found that strongly locomotion-modulated cell clusters during visual stimulation were aligned with M2− interpatches, while weakly modulated cells clustered under M2+ patches. Unlike M2+ patch cells, pairs of M2− interpatch cells showed increased correlated variability of calcium transients when the sites in the visuotopic map were far apart, suggesting that activity is integrated across large parts of the visual field. Pathway tracing further suggests that strong locomotion modulation in L2/3 M2− interpatch cells of V1 relies on looped, like-to-like networks between apical dendrites of MOs-, PM- and RSP-projecting neurons and feedback input from these areas to L1. M2− interpatches receive strong inputs from SST neurons, suggesting that during locomotion these interneurons influence the firing of specific subnetworks by controlling the excitability of apical dendrites in M2− interpatches.

**eLife digest:** During body, head and eye movements, many cells in the visual system increase their firing rate in order to distinguish between self-generated and externally generated image motion. While progress has been made in determining the brain circuits and cell types underlying this state-dependent signal, it is not known whether movement-modulated cells in primary visual cortex (V1) are distributed randomly or are spatially clustered. ***Meier et al.,*** found that locomotion modulation is greater in periodically spaced clusters of layer (L) 2/3, which are aligned with M2 muscarinic acetylcholine receptor-negative interpatches of L1. Cells aligned with interpatches have higher sensitivity to the direction of visual motion at high speed (***Ji et al., 2015***). Cell pairs between visuotopically distant interpatch clusters show increased response correlations, suggesting signal integration across large parts of the visual field. The results further demonstrate that L1 M2− interpatches are densely innervated by somatostatin (SST)-expressing neurons, are targets of apical dendrites of MOs-(secondary motor cortex), PM-(posteromedial area) and RSP-(posterior retrosplenial cortex)-projecting neurons and are recipients of inputs from these sources. This indicates that L1 interpatches in V1 are targets for non-retinal movement-related feedback signals.

## Introduction

Sensory responses are not rigid representations of stimuli formed in the brain disconnected from the body, but are shaped by actions in the real world (***Bermejo et al., 2020***). A prominent example of such effects were found in V1, in which locomotion increases the gain of visual responses, and running speed drives neuronal activity even in the dark (***Niell and Stryker, 2010; Pakan et al., 2016***; ***Saleem et al., 2013***). This increase in responsiveness has been shown to improve the fidelity of stimulus representations across the neuronal population (***Dadarlat and Stryker, 2017***) and enhance visual detection (***Bennett et al., 2013***). Reliable interactions between inputs carrying information about the speed of locomotion and visual flow are important for computing an animal’s location during navigation through the environment (***Chen et al. 2013***). Such interactions are driven by inputs from multiple sources, including speed-sensitive neurons in the mesencephalic locomotor region (***Carvalho et al., 2020***; ***Lee et al., 2014***) and release of inhibition by SST neurons in V1 (***Dipoppa et al., 2018; Fu et al., 2014)*.**

Neural activity of visual cortex has also been shown to be modulated by orienting movements of the head and visually guided movements of the eyes (***Bouvier et al., 2020; Itokazu et al., 2018; Parker et al., 2022***; ***Velez-Fort et al., 2018***). Unlike the proprioceptive inputs elicited by locomotion, the sensory inputs from head movements derive mostly from the vestibular organ (***Bouvier et al., 2020***), whose signals reach V1via the retrosplenial cortex (***Vélez-Fort et al., 2018***). Recordings in freely moving rats suggest that head movement signals also originate in secondary motor cortex (MOs) (***Guitchounts et al., 2020***), which sends efference copy signals to V1 (***Leinweber et al., 2017***) that are required for guiding voluntary eye movements (***Itokazu et al., 2018)*.**

Locomotion also affects contextual visual processing, by reducing the suppression of responses in the receptive field (RF) center from the RF surround (***Ayaz et al., 2013***). Such influences have been shown to depend on inhibition by SST-expressing GABAergic neurons (***Adesnik et al., 2012***). A similar dependency on SST-mediated inhibition has been reported for head movement-related vestibular signals recorded in V1 (***Bouvier et al., 2020***).

Although widespread, the locomotion-related proprioceptive, vestibular and motor efference copy signals are highly variable and are expressed by less than half of V1 neurons (***Ayaz et al., 2013; Erisken et al., 2014; Saleem et al., 2013***). This suggests that the motor-related modulations of V1 activity are used in specific subnetworks, specialized for differentiating between self-generated visual motion and external object motion during navigation and rapid eye movements (***Fajen and Matthis, 2013***; ***Miura and Scanziani, 2022***). Such functionally distinct subnetworks are a prominent, distributed feature of mouse visual cortex (***Sit and Goard, 2020***).

Here we investigated whether there are distinct subnetworks within V1, differentially linked to the M2+ patch/M2-interpatch system (***Ji et al., 2015***), in which visual responses are preferentially modulated by locomotion. The results show that locomotion modulation is stronger in M2− interpatches than in M2+ patches, and that visual responses during running in M2− interpatch cells have increased pairwise correlations in trial-to-trial variability when spaced far apart at distant sites of the visuotopic map. Further, we found that L1 M2− interpatches overlay with inputs from local SST interneurons, long-range inputs from MOs, PM and RSP and apical dendritic tufts of neurons which project to these targets. These findings suggest that locomotion modulation is a differentiating feature of M2− interpatch cells, associated with a distinct subnetwork of the medial dorsal stream (***Wang et al., 2012***).

## Results

### Locomotion modulation of visual responses is more robust in M2− interpatch cells

We injected AAV2/1.hSynapsin.Flex.GCaMP6f at multiple sites of V1 of adult Chrm2-tdTomato knock-in mice crossed with Emx1-Cre conspecifics. The progeny expressed M2 muscarinic acetylcholine receptor-tdTomato fusion protein in patterns tightly overlapping with the endogenous *Chrm2* gene. Cell type-specific transduction of GCaMP6f allowed the comparison of calcium responses from L2/3 pyramidal cells (PCs) aligned with M2+ patches or M2− interpatches in L1. A head plate was implanted over a cranial window for access to the injection sites (***Figure 1a***). Neural activity of L2/3 PCs was recorded as calcium responses in the tangential plane, using 2-photon microscopy. Awake mice were head-fixed, placed on a wheel, and allowed to run spontaneously during three periods with different visual stimulation conditions (***Figure 1b***). First, a circular drifting grating (20° in diameter) was presented on a monitor at 24 different locations, and responses were used to compute the average RF of hundreds of cells. Second, the screen was turned off for 10 minutes, during which time spontaneous neural activity was recorded in total darkness (< 0.1 cd/m^2^). Third, a drifting circular square-wave grating (30° in diameter) was presented with a range of spatial frequencies (SF; 0.01–1.6 c/°), temporal frequencies (TF; 0.1-12Hz), and orientations (OS; 51° increments). After recording four to five sessions per mouse at separate non-overlapping locations of V1, mice (N = 9) were sacrificed.

**Figure 1.**
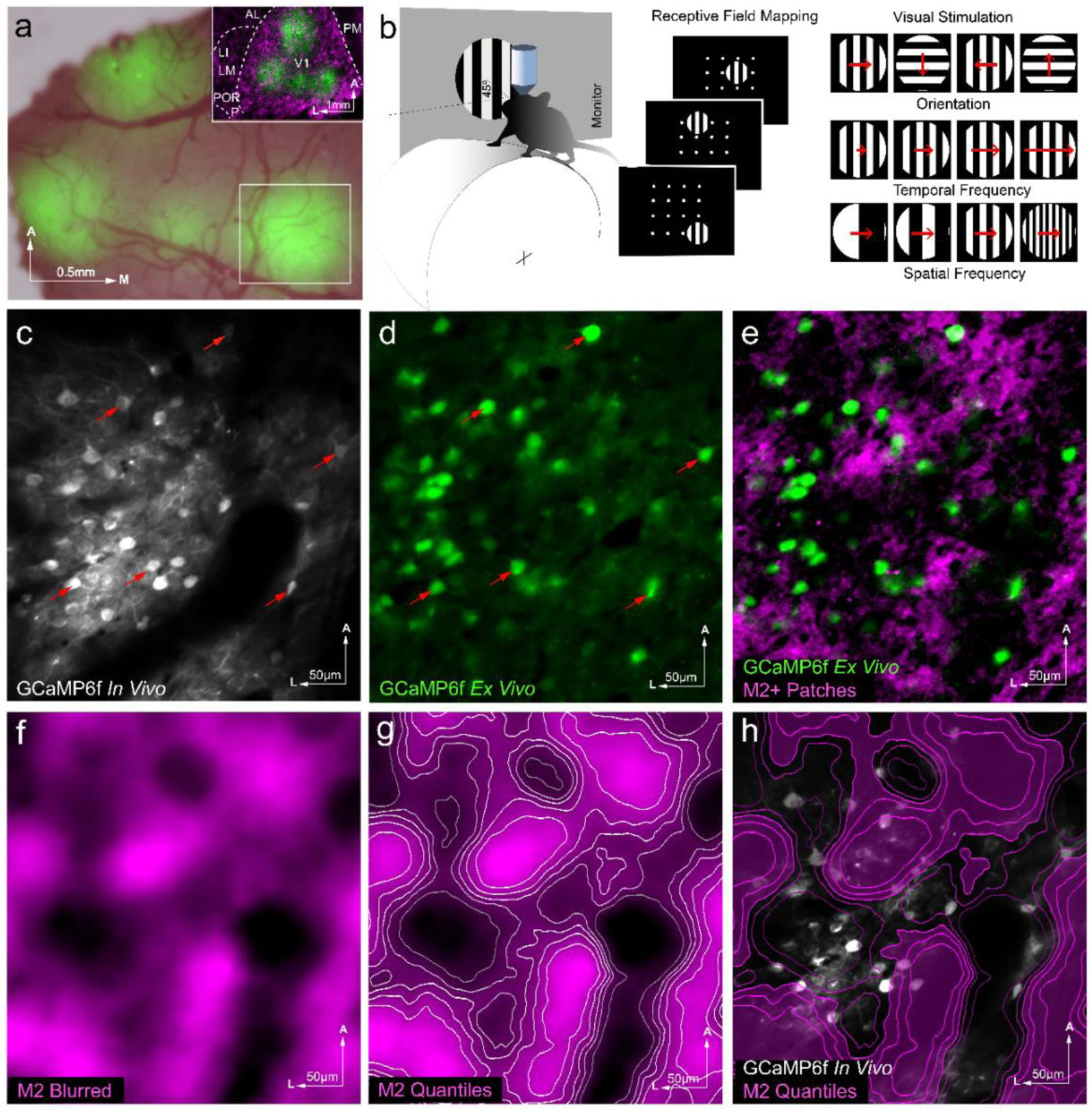
Experimental setup, visual stimulation, and tissue alignment protocol for calcium imaging in V1 of Chrm2tdT x Emx 1-Cre mice. (**a)** *In vivo* image of four injection sites of AAV2/1.hSyn.Flex.GCaMP6f in V1 after head plate implantation, visualized by overlaying brightfield image of surface blood vessels with GFP expression. Inset shows *ex-vivo* tangential section with GCaMP6f injection sites (green) overlaid with Chrm2tdT expression in L1 (magenta). AL (anterolateral area), LI (laterointermediate area), LM (lateromedial area), P (posterior area), PM (posteromedial area), POR (postrhinal area), V1 (primary visual area). (**b**) Schematic of two photon calcium recording in V1 of awake headfixed mice running on a wheel, facing a drifting square wave grating stimulus on a monitor (left). Mapping of receptive fields by stimulating with 20°-wide patches of a drifting gratings at one of 24 locations (center). Stimuli used for recording tuning curves to square wave gratings of varying spatial frequencies, temporal frequencies, and orientations (right). The orientation and length of red arrows indicate the direction and speed of stimulus motion. (**c-h**) Alignment of recorded cells with M2+ patches. (**c**) Time-averaged *in vivo* GCaMP6f signals from pyramidal cells showing somas of active V1 L2/3 cells contained in the white box outlined in panel **a**. Examples of responsive cells are labeled by arrows. (**d**) Cellular GCaMP6f expression (arrows) in *ex vivo* paraformaldehyde-fixed 40 µm tangential section at the same location as shown in panel **c**. (**e**) Overlay of *ex vivo* GCaMP6f expression shown in panel **d** aligned with *ex vivo* M2+ patches in L1. (**f**) Image of M2+ patches shown in panel **e** after high-pass filtering and blurring. (**g**) Image from panel **f** with contours outlining 6 intensity quantiles. (**h**) *In vivo* GCaMP6f image from panel **c** overlaid with M2 quantiles from panel **g**, allowing cells (white) from *in vivo* recordings to be assigned to M2 quantiles. M2+ patches are colored magenta.

While *in vivo* imaging of tdT expression was readily achieved, the periodic pattern in L1 was often distorted by surface blood vessels, which disguised the periodic M2 expression pattern and hindered identification of M2+ patches and M2− interpatches. To get around this problem, mice were perfused with fixative, the cortex was flatmounted and sectioned in the tangential plane. The *ex vivo* Chrm2tdT expression pattern in L1 was imaged under a fluorescence microscope and aligned with the M2 pattern imaged *in vivo* (see, Methods). Although *ex vivo* and *in vivo* images lined up well across the 330 x 330 um regions of interest, registration was further optimized by global warping. M2 images were high-pass filtered, blurred, and divided into six intensity quantiles (***Figure 1f, g***). The top 3 quantiles were considered to be M2+ patches and the bottom 3 were considered to be M2− interpatches. *In vivo* recorded cells (***Figure 1c***) were recovered in fixed sections by the green fluorescence of Ca^2+^-bound GCaM6f (***Figure 1d***). Responsive cells were aligned with their counterparts in *ex vivo* sections and registered to the M2 pattern (***Figure 1e, h***), using matching patterns of vertically descending blood vessels and labeled cells as fiducial markers for warping. The warping function’s effect was mostly rotation of the image. When present, stretching was negligible and confined to the margins of the image (***Figure 1 c, d*; *Figure 1 – Supplement 1***).

For each cell, a locomotion modulation index (LMI) (***Pakan et al., 2016*)** was computed as (R_L_ – R_S_)/(R_L_ + R_S_), where R_L_ is the mean ΔF/F stimulus response during locomotion trials (≥ 1cm/s, and R_S_ is the mean response during immobility (< 1cm/s). ***Figure 2a*** shows SF, TF, and OS tuning curves computed during rest (black curves) and locomotion (red curves) of example cells near the center of M2− interpatches (quantiles 1-2), the vicinity of M2+ patch centers (quantiles 5-6), or in the transition zone between M2+ and M2− clusters (quantiles 3-4). The inset in ***Figure 2a*** shows the stimulus response time course of an example M2− interpatch cell to its preferred SF, for which the response during locomotion trials (n = 4 trials) was approximately twice its response to stationary trials (n = 6 trials). Each cell was classified as either locomotion-tuned or not locomotion-tuned, by determining whether its visual responses were of significantly different magnitude in locomotion and stationary trials (unpaired t-test, α = 0.05). Most locomotion-tuned cells (89.3%, 528/591) showed a positive gain to visual stimulation with a mean LMI of 0.19 ± 0.01. M2− interpatch cells showed a 44.6 ± 12% greater LMI than M2+ patch cells (***Figure 2b***; N = 9 mice; n = 1077 cells; r = −0.08; *p* < 0.01, Pearson correlation).

**Figure 2.**
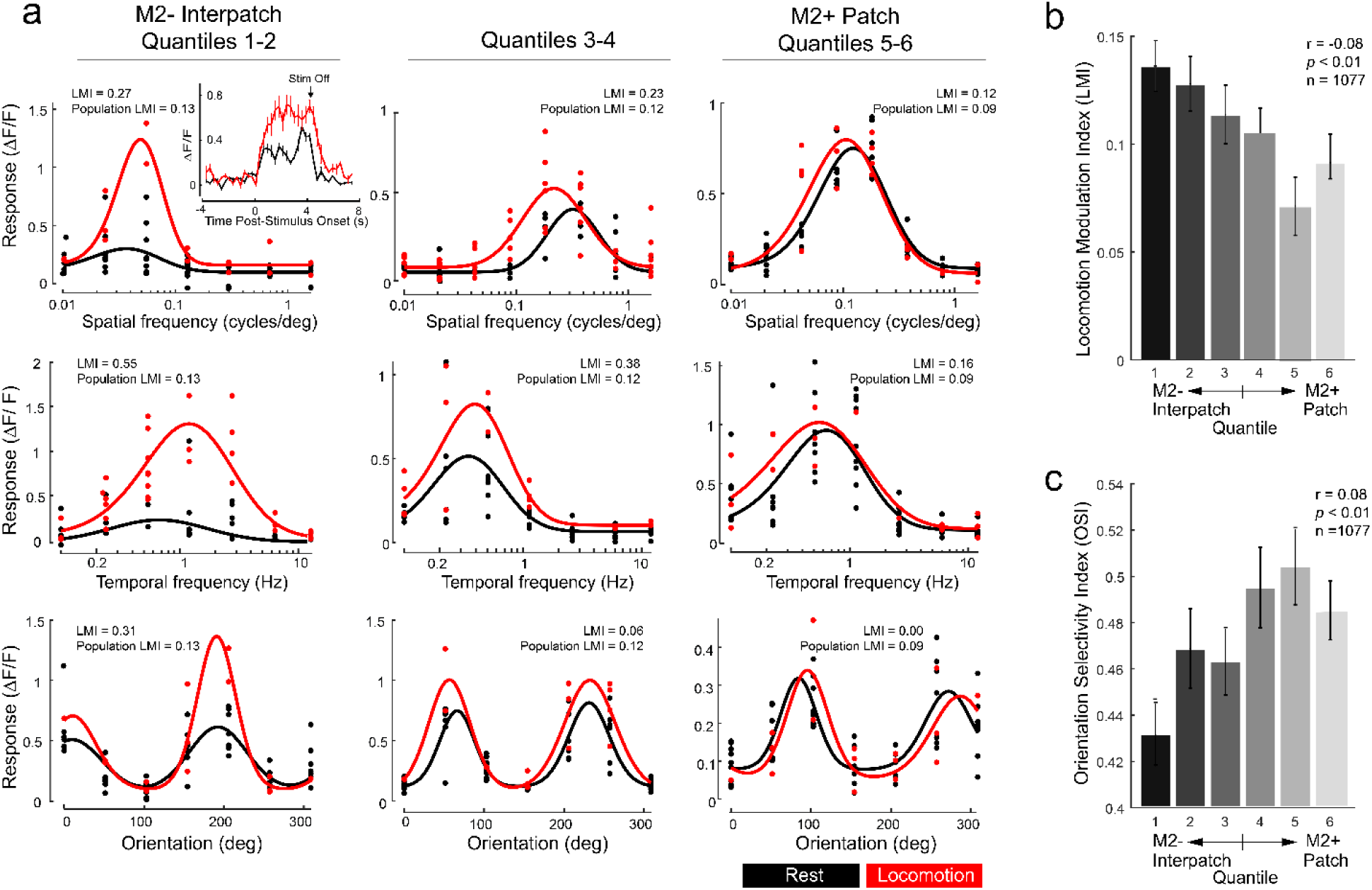
Locomotion modulation and orientation tuning in L2/3 pyramidal cells of V1, aligned with M2+ patches and M2− interpatches of L1. (**a**) Tuning curves from example cells for spatiotemporal stimulus parameters. Red traces show tuning curves generated by trials with locomotion (≥ 1 cm/sec). Black curves indicate trials during which mice were immobile (< 1 cm/sec). The leftmost column contains M2− interpatch cells (quantiles 1-2), the rightmost contains M2+ patch cells (quantiles 5-6), the middle column represents cells at the border between M2+ patches and M2− interpatches (quantiles 3-4). Fitted tuning curves are shown with dots indicating responses from individual trials. Inset: trial-averaged time courses of a sample M2− interpatch neuron to its preferred spatial frequency stimulus during locomotion (red; n = 4 trials) and rest (black; n = 6 trials). Error bars show SEM in each time bin. (**b**) Locomotion Modulation Index (LMI) of visual responses in M2+ patch and M2− interpatch cells shows preferential modulation of M2− interpatch cells. (**c**) Distribution of Orientation Selectivity Index (OSI) computed from combined trials during locomotion and immobility shows increased selectivity of M2+ patch cells. Error bars represent SEM across cells.

Because experience can shape visual responses and their locomotion modulation in mouse V1 (***Hoy and Niell, 2015; Kaneko and Stryker, 2014; Wang et al., 2024***), we sought to rule out the possibility that our results were affected by LMI systematically varying with the mouse’s experience across the sessions. We found that LMI of individual cells did not correlate with the number of prior sessions undergone (r = 0.006, *p* = 0.882, Pearson correlation) (***Figure 2 – Supplement 2.1a).*** To further test whether LMI is independent of task repetition and visual stimulation we compared a set of linear models of LMI containing or omitting these predictors. These included a model of LMI predicted by session number and a model of LMI predicted by both session number and module, i.e. M2+ patch vs. M2− interpatch. We found that a model using only M2 module as a predictor was a superior fit to the data (Akaike information criterion, AIC = −477.3) than either a model using session number (AIC = −467.4) or session number and module (AIC = −475.4; ***Figure 2 – Supplement 2.1b***). These results indicate that LMI did not vary systematically with task experience. Thus, session number was ignored for all subsequent analyses.

### Orientation selectivity is stronger in M2+ patches

Previous electrophysiological recordings in V1 of anesthetized mice found that L2/3 cells aligned with M2+ patches are specialized for processing the orientation of stimulus contours (***Ji et al., 2015***), raising the question of whether similar selectivities can be observed in awake mice. The strength of orientation tuning was measured by the orientation selectivity index (OSI) (***Mazurek et al., 2014),*** computed as (R_pref_ *−* R_ortho_)*/*(R_pref_ *+* R_ortho_), where R_pref_ is the average ΔF/F response of the cell to its preferred orientation, and R_ortho_ is its response to orientations tilted 90° away from the preferred orientation. We found that OSI was higher in M2+ patches than M2− interpatches, indicating that M2+ patch cells were more strongly tuned for orientation than M2− interpatch cells (***Figure 2c***; n = 1077 cells; r = 0.08, *p* < 0.01, Pearson correlation). M2+ patches and M2− interpatches showed similar half-width at half-maximum (HWHM) responses for SF, TF, and orientation (***Figure 2 – Supplement 2.2a-f***). Similarly, we found no difference in peak SF and TF between M2+ patches and M2− interpatches (***Figure 2 – Supplement 2.2g, h).*** Thus, locomotion did not impact the average tuning width and peak sensitivity of cells to the different types and stimulus parameters used.

### M2+ patch pairs show increased activity correlations in darkness

M2+ patches and M2− interpatches of L1 have been shown to belong to distinct networks (***Burkhalter et al., 2024***; ***D’Souza et al., 2019; Ji et al., 2015; Meier et al., 2021***). This raised the question of whether the cells in the layers below L1 M2+ patches and M2− interpatches have discrete pairwise activity correlations (see Methods), which may facilitate the flow of information through the network. Spontaneous correlations were computed as the Pearson correlation coefficient between the ΔF/F traces of two cells throughout the dark period, during both stationary and locomotion epochs. Cell pairs were grouped into M2+/M2+, M2−/M2− and mixed M2+/M2− pairs. Previous studies in mouse V1 found that pairwise correlations decrease with increasing distance between cells (***Denman and Contreras, 2014; Yu et al., 2019***). To analyze this, we grouped cell pairs by distance, using 25µm-wide bins. We first examined pairwise ΔF/F Pearson correlation coefficients during the 10-minute dark period (temporal bin size 0.33 s). We found a striking difference between pair types: in every cell-pair distance bin from 0-275µm, M2+/M2+ pairs were significantly more correlated than M2+/M2− and M2−/M2− pairs (***Figure 3a***). This difference between pair types was absent in pairs that were farther apart than ≥ 275µm. When analyzing cell pairs without sorting by distance, M2+/M2+ pairs also showed greater activity correlations than M2+/M2− and M2−/M2− pairs (n = 74,593 pairs, N = 9 mice, ANOVA, *p* < 10^-10^).

**Figure 3.**
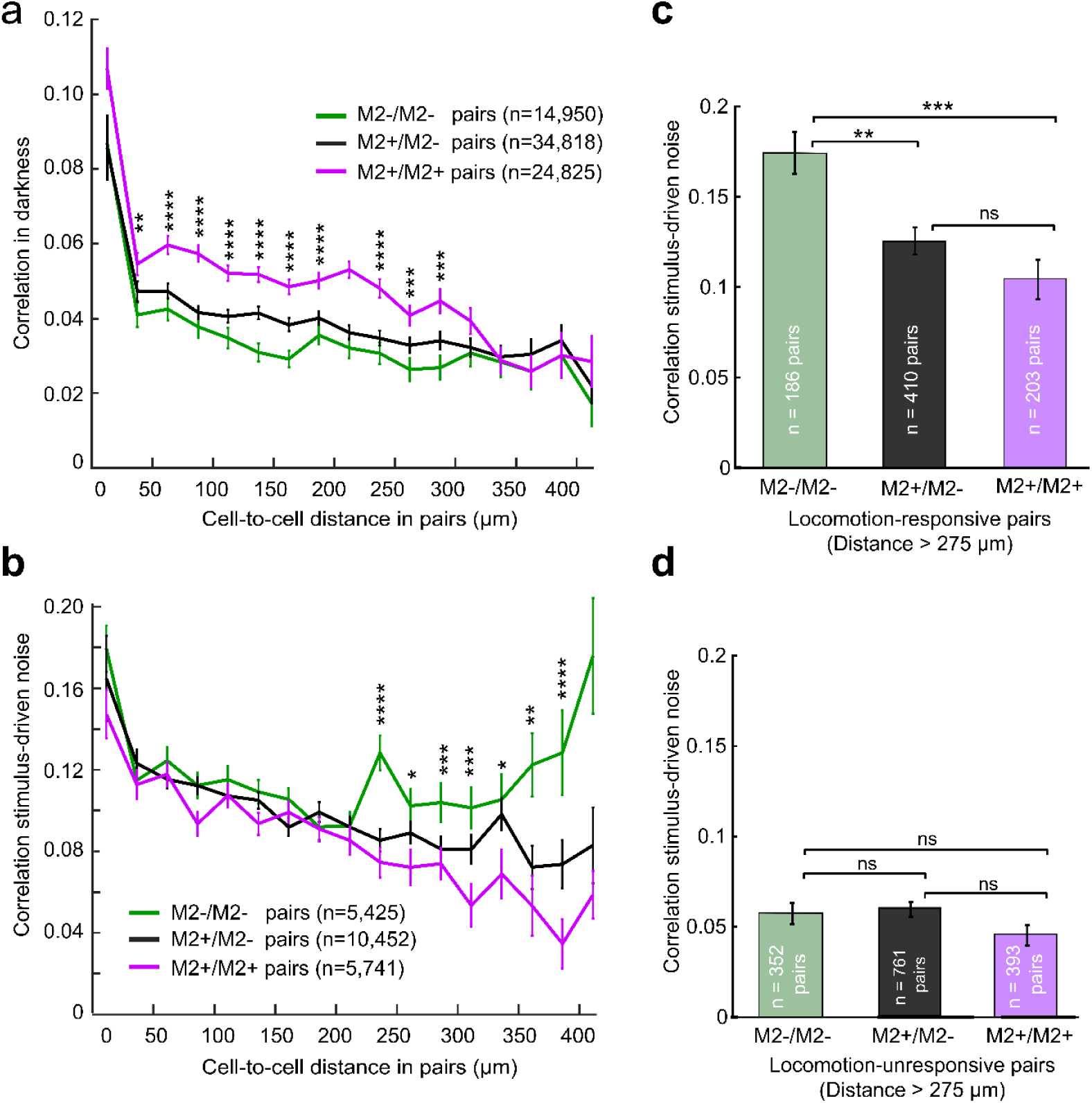
Pairwise correlated activity of L2/3 M2+ patch and M2− interpatch cells in V1. Pairs were divided into those containing two M2+ patch cells (M2+/M2+, magenta), two M2− interpatch cells (M2−/M2−, green), and one of each type (M2+/M2−, black). (**a**) Pearson correlation coefficient of ΔF/F signals (mean ± SEM) of cell pairs in darkness at different cell-to-cell distances. Asterisks indicate significant differences between correlation coefficients in M2+/M2+ of M2−/M2− and M2+/M2− cell pairs at different cell-to-cell distances (t-test; ⁎ = *p* < 0.05, ⁎⁎ = *p* < 10^-2^, ⁎⁎⁎ = *p* < 10^-3^, ⁎⁎⁎⁎ = *p* < 10^-4^). (**b**) Coefficient of response noise correlation (mean ± SEM) between cell pairs during visual stimulation at different cell-to-cell distances. Asterisks same as in panel **a**. (**c**) Comparison of response noise correlations (mean ± SEM) between cell pairs in which both cells were locomotion-responsive and the cell-to-cell distance was > 275 µm. Asterisks indicate significance level (t-test with Bonferroni correction; ns = not significant, ⁎ = *p* < 10^-2^, ⁎⁎ = *p* < 10^-3^, ⁎⁎⁎ = *p* < 10^-4^). (**d**) Comparison of response noise correlations (mean ± SEM) between cell pairs in which neither cell was locomotion-responsive and the cell-to-cell distance was > 275µm. Note, that the number of pairs in panel **d** is larger than in **c**, which includes pairs with one locomotion-tuned and one non-locomotion-tuned cell. Asterisks same as in panel **c.**

### Responses of widely separated M2−/M2− cell pairs are more correlated than distant M2+/M2+ and M2+/M2− cell pairs

We next computed the correlated variability of visual responses, sorting pairs by M2+ patch and M2− interpatch module and distance. Response noise correlations were computed by finding the Pearson correlation between response magnitudes in every trial with identical spatiotemporal features (SF, TF, and orientation) of visual stimulation, then averaging this value across all unique stimuli (***Montijn et al., 2014***). We found that unlike activity correlations in darkness, visual response noise correlations showed almost no module-specific differences in bins from 0 - 275 µm (***Figure 3b***; n = 21,595 pairs, N = 9 mice). Notably, however, cell pair noise correlations diverged by module type when farther apart. At longer (> 275 μm) distances M2−/M2− interpatch pairs (n = 5,429) showed increased correlation values, whereas for M2+/M2+ patch (n = 5,741) and mixed M2+/M2− pairs (n = 10,425) correlations continued to decrease as the distance between cells increased (***Figure 3b***).

Earlier findings that running speed affects pairwise spike correlations (***Erisken et al., 2014***) raised the question of whether stimulus-driven response noise correlations of cell pairs were mediated by locomotion. Pairs were divided into those in which both cells were locomotion-tuned (see above) and pairs in which neither were locomotion-tuned. Pairs with one locomotion-tuned and one non-locomotion-tuned cell were not included in this analysis. We found that for topographically distant (> 275 µm) locomotion-tuned pairs in V1, pair type with respect to M2 module played a significant role in response noise correlations (***Figure 3c***), with M2−/M2− pairs (n = 186) having 67% higher correlation coefficients than M2+/M2+ pairs (n = 203; *p* < 10^-4^, t-test with Bonferroni correction) and 39% higher correlation coefficients than mixed M2+/M2− pairs (n = 410; *p* < 10^-3^). Widely separated pairs whose responses were not locomotion-tuned showed no difference between M2−/M2− (n = 352) vs. M2+/M2+ pairs (n = 393; *p* = 0.24), M2−/M2− vs. M2+/M2− pairs (n = 761, *p* = 0.13), or M2+/M2+ vs. M2+/M2− pairs (*p* = 0.74) (***Figure 3d***). These results suggests that locomotion selectively affects the correlated activity of M2− interpatch cells pairs which are farther apart than 275 µm.

### Looped module-specific like-to-like connections between V1 and higher cortical areas including the lateral posterior thalamic nucleus

Locomotion signals may originate from MOs, which has been shown to provide motor efference copy feedback signals to V1 (***Leinweber et al., 2017***). To study the architecture of the underlying network, we injected a 1:1 mixture of anterograde AAV2/1.hSyn.EGFP.WPRE.bGH and retrograde AAV2retro.CAG.Cre into MOs of Ai9 mice (***Figure 4a, b***). Images of M2 immunostained sections through flatmounted cortex were overlaid with anterogradely labeled MOs→V1 projections to L1, to determine the density of MOs input to M2+ patches and M2− interpatches. We found that MOs axonal projections to V1 were non-randomly distributed, with significantly (Pearson correlation, r = −0.41, *p* = 0.02, N = 5 mice) denser terminal branching in M2− interpatches (***Figure 4c-e, j***). Notably, retrogradely labeled apical dendrites of V1→MOs-projecting neurons in L1 (***Figure 4f***) showed a similarly strong preference for M2− interpatches (Pearson correlation, r = −0.62, *p* = 4.9 × 10^-5^, N = 6) (***Figure 4g-i, k***).

**Figure 4.**
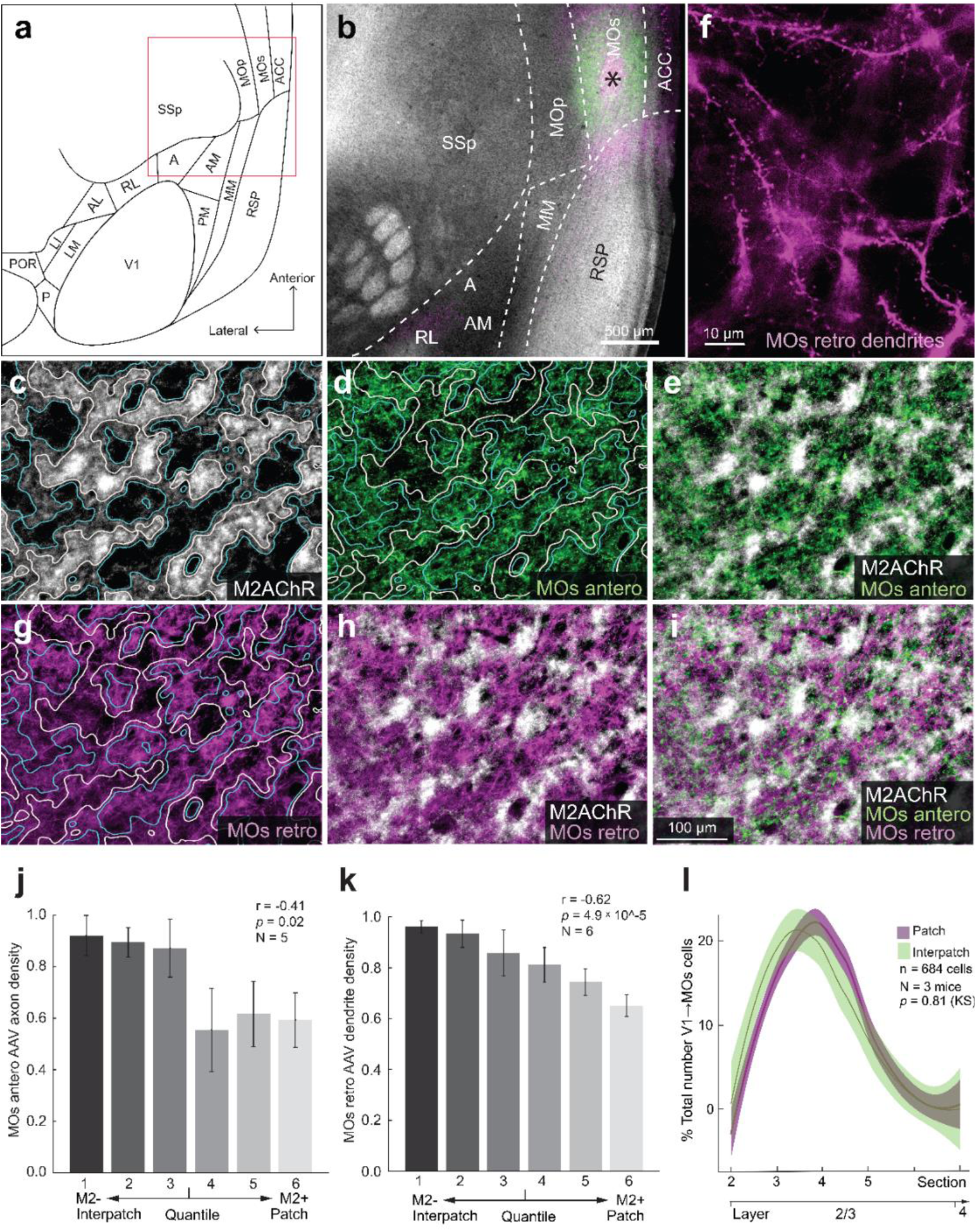
Looped modular like-to-like of connections between V1 and secondary motor area (MOs. **(a**) Diagram of flatmap of left mouse cerebral cortex. Boxed area indicates location of image shown in panel **b**. A (Anterior area), ACC (Anterior cingulate cortex), AL (Anterolateral area), AM (Anteromedial area), LI (Lateral intermediate area), LM (Lateromedial area), MM (Mediomedial area), MOp (Primary motor area), MOs, P (Posterior area), PM (Posteromedial area), POR (Postrhinal area), RL (Rostrolateral area), RSP (Posterior retrosplenial area), SSp (Primary somatosensory area), V1 (Primary visual area). (**b**) Tangential section through L4 of flatmounted cortex of Ai9 mouse, stained with an antibody against M2AChR (M2) (white). Injection site in MOs (asterisk) of cocktail of viral tracers (AAV2/1-hSyn-EGFP.WPRE.bGH [green], rAAV2-Retro-CAG.Cre [magenta]). (**c**) Tangential section through L1 of flatmounted V1. White regions are M2AChR immunostained M2+ patches, outlined by white lines. Dark regions that lack M2, represent M2−interpatches, and are outlined by blue lines. (**d**) Anterogradely AAV-labeled MOs→V1 projections (green) in L1. (**e**) Overlay of panels **c** and **d**, showing preferential input of MOs→V1 axons to M2− interpatches. (**f**) Retrogradely labeled spine-covered apical dendrites of V1→MOs projecting cells arborizing in L1 of V1. (**g**) Apical dendrites of retrogradely AAV-labeled V1→MOs projecting neurons in L1 (magenta) preferentially branch in M2− interpatches outlined by white lines. M2+ patches are surrounded by blue lines. (**h**) Overlay of apical dendrites (magenta) and M2+ patches (white). (**i**) Overlay of MOs→V1 axon projections (green), apical dendrites of V1→MOs projection neurons (magenta) and M2+ patches (white) in L1 of V1. (**j**) Labeling density of MOs→V1 axons in different M2 quantiles. Pearson correlation (r), error bars ±SEM. (**k**) Labeling density of dendrites of V1→MOs projecting cells different M2 quantiles. Pearson correlation (r), error bars ±SEM. (**l**) Laminar distribution of retrogradely labeled cells bodies shows that most cells are located in L2/3 and exhibit no preferential alignment with M2+ patches (magenta) and M2− interpatches (green). KS test. Shading ±SEM.

Next, we investigated whether module-selective loops, shown by the overlapping axons and dendrites in the V1↔ MOs pathway with L1 (***Figure 4c-k***), are general features by which V1 interacts with cortical areas and the thalamus. We injected a mixture of AAV2retro.CAG.Cre / AAV2/1.hSyn.EGFP.WPRE.bGH into LM, RL, PM, RSP and the lateral posterior nucleus (LP) of the thalamus. We then aligned anterogradely labeled axons and retrogradely labeled cells including dendrites with M2 immunolabeling in L1 of V1. We found that anterogradely labeled feedback projections from each of the higher visual areas were strikingly clustered: LM→V1 (***Figure 5a-c, g***) and RL→V1 (***Figure 5 – Supplement 1 a-c, g***) preferentially terminated in M2+ patches, whereas PM→V1 (***Figure 6a-c, g***) and RSP→V1 inputs (***Figure 6, Supplement 1a-c, g***) preferred M2− interpatches. In each case, retrogradely labeled feedforward-projecting cell bodies in V1 were found in all layers, with a strong bias for L2/3 (***Harris et al., 2019; Young et al., 2021***).

**Figure 5.**
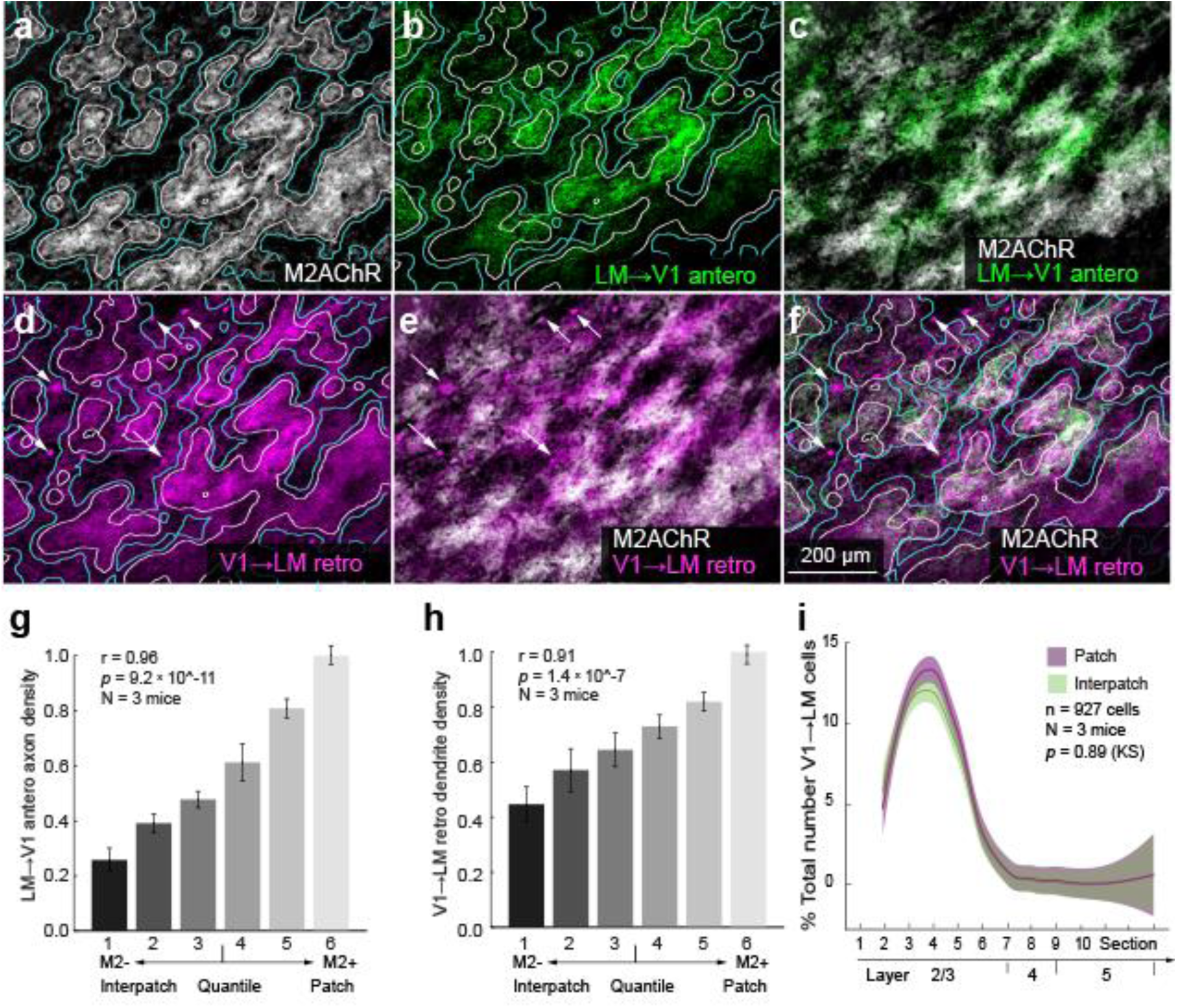
Looped clustered like-to-like connections between V1 and LM. (**a**) Tangential section through L1 of V1 of Ai9 mouse stained with an antibody against M2AChR (M2). M2+ patches (white) are outlined by white lines, M2− interpatches are indicated by blue lines. (**b)** Anterogradely AAV-labeled (for tracer, see Figure 4) LM→V1 axons (green) in L1 of V1. (**c**) Overlay of panels **a** and **b**. Axons preferentially terminate in M2+ patches. (**d)** Apical dendrites of retrogradely AAV-labeled V1→LM-projecting neurons (magenta) in L1 outlined by white lines. M2− interpatches outlined by blue lines. Note that the section is not perfectly parallel to the pial surface. It runs from right to left through L1 and the top of L2, which shows cell bodies (arrows) in M2+ patches and M2− interpatches. (**e**) Apical dendrites of V1→LM-projecting neurons (magenta) preferentially branch in M2+ patches (white). Retrogradely AAV-labeled cell bodies (arrows). (**f**) Overlay of panels **d** and **e**. Apical dendrites of V1→LM-projecting neurons (magenta) terminate in M2+ patches (white). Retrogradely AAV-labeled cell bodies (arrows). (**g**) LM→V1 axon projection density in different M2 quantiles of L1. Pearson correlation (r), error bars ±SEM. (**h**) Labeling density of retrogradely AAV-labeled V1→LM dendrites in different M2 quantiles of L1. Pearson correlation (r), error bars ±SEM. (**i**) Laminar distribution of retrogradely labeled V1→LM-projecting cells shows that most cell bodies are located in L2/3 and are randomly distributed relative to M2+ patches (magenta) and M2− interpatches (green) in L1. KS test. Shading ±SEM.

**Figure 6.**
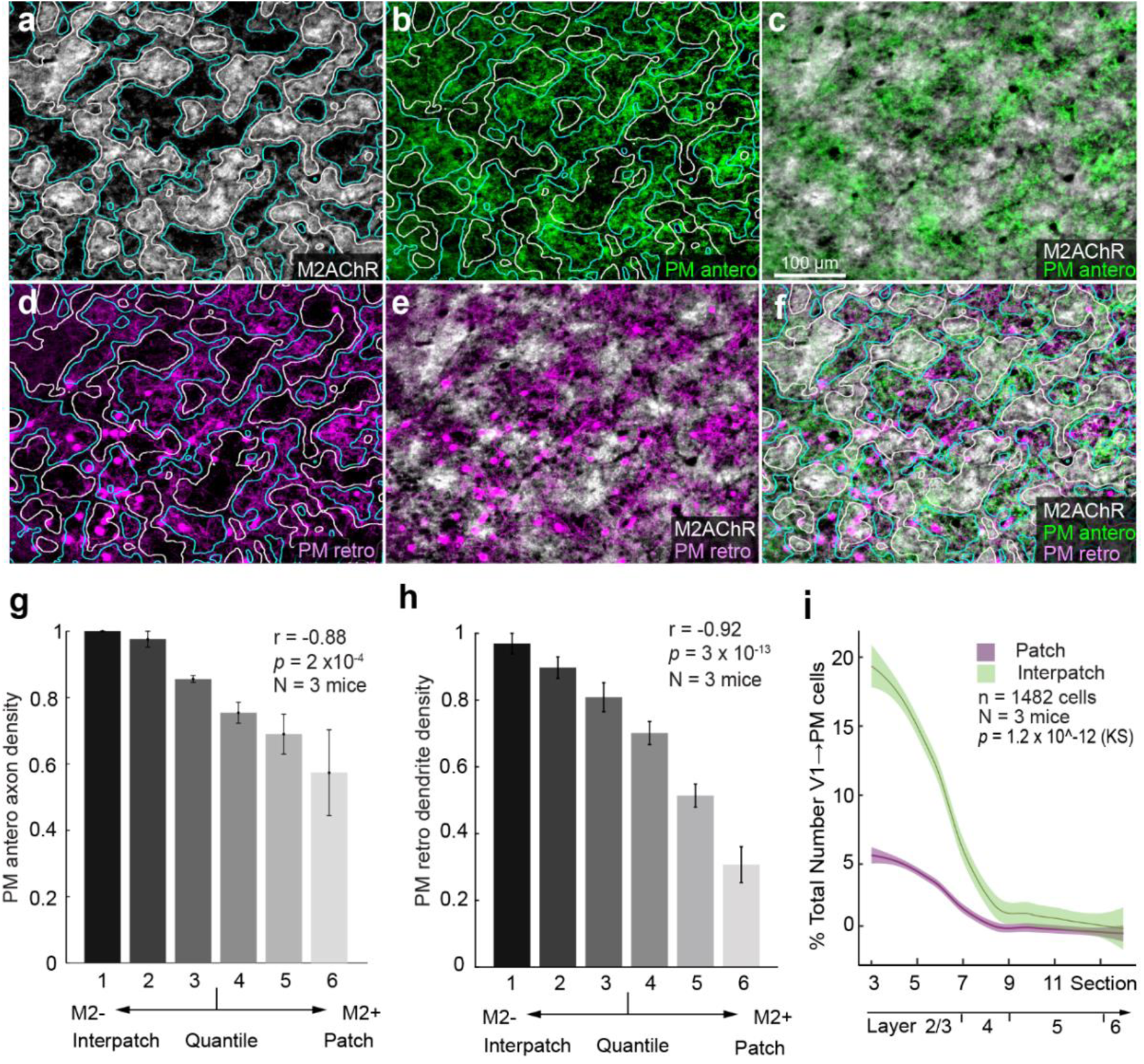
Looped modular like-to-like of connections between V1 and PM. (**a**) Tangential section through L1 of V1 of Ai9 mouse stained with an antibody against M2AChR (M2). M2+ patches (white) are outlined by white lines, M2− interpatches by blue lines. (**b**) Anterogradely AAV-labeled PM→V1 axon projections (green) in L1 of V1 terminating in M2− interpatches outlined by blue lines. M2+ patches outlined by white lines. (**c**) Interdigitation of PM→V1 axon projections (green) and M2+ patches (white). (**d**) Apical dendrites of retrogradely AAV-labeled V1→PM-projecting cell bodies (magenta) at the L1/2 border of V1, preferentially arborize in M2− interpatches (blue outlines) of L1. M2+ patches are outlined by white lines. (**e**) Interdigitation of dendrites of retrogradely labeled V1→PM-projecting cells (magenta) and M2+ patches (white). (**f**) Overlay of panel **e** with anterogradely labeled axons shown in panel **c**. (**g**) Density of PM→V1 axon projections in different M2 quantiles of L1. Pearson correlation (r), error bars ±SEM. (**h**) Labeling density of retrogradely labeled dendrites of V1→PM-projecting neurons in L2/3 aligned with different M2 quantiles of L1. Pearson correlation (r), error bars ±SEM. (**i**) Laminar distribution of retrogradely labeled cells bodies shows that most cells are preferentially aligned with M2− interpatches (green) in L1. Magenta indicates cells aligned with M2+ patches. KS test. Shading ±SEM.

Notably, in each pathway retrogradely labeled apical dendrites in L1 were non-randomly distributed: V1→LM (***Figure 5d-f, h***) and V1→RL were clustered in M2+ patches (***Figure 5, Supplement 1 c, d-f, h),*** while V1→PM (***Figure 6 d-f, h***), V1→ RSP (***Figure 6, Supplement 1d-f, h***) and V1→ LP were significantly denser in M2− interpatches (***Figure 6, Supplement 2a-e***). These overlapping patterns suggest that V1 forms looped like-to-like connections with distinct preferences for either M2+ patches (LM, RL) or M2− interpatches (MOs, PM, RSP, LP).

### Diverse distributions of cell bodies

The module-specific overlap of axons and dendrites in L1 of V1 raised the question of whether the parent cell bodies in the layers below are vertically aligned with the corresponding M2+ patches or M2− interpatches. Unexpectedly, we found random tangential distributions in some pathways and clusters in others. The majority of V1→MOs-, V1→LM-, V1→AL-(not shown), V1→RL-, V1→PM- and V1→RSP-projecting neurons were located in L2/3 (***Figure 4l, Figure 5i, Figure 5 Supplement 1i, Figure 6i, Figure 6 Supplement 1i),*** except for the V1→LP-projecting cells located in deep L5 (***Figure 6 Supplement 2e***). Unlike the clustering of dendrites in M2− interpatches (***Figure 4k***) the parent cell bodies of V1→MOs-projecting neurons were randomly distributed (n = 684 cells, N = 3 mice, *p* = 0.81, Kolmogorov-Smirnov [KS] test) (***Figure 4l***). Similarly, we found randomly distributed cell bodies in the V1→LM (n = 927 cells, N = 3 mice, *p* = 0.89, KS), V1→AL (n = 756 cells, N = 2 mice, *p* = 0.27, KS) (not shown) and V1→RL (n = 1515 cells, N = 3 mice, *p* = 0.08, KS) pathways, in which the dendrites were clustered in M2+ patches (***Figure 5h, i; Figure 5 Supplement 1h, i***). In sharp contrast, in V1→PM (n = 1482 cells, N = 3 mice, *p* =1.2 × 10^-12^, KS), V1→RSP-(n = 1453 cells, N = 3 mice, *p* = 7 × 10^-9^, KS) and V1→LP-projecting (n = 803, N = 3 mice, *p* = 0.02, N = 3, KS) pathways, cell bodies were preferentially aligned with M2− interpatches (***Figure 6i, Figure 6 Supplement 1i, Figure 6 Supplement 2e***). These results show that while in each pathway axons and dendrites are preferentially targeted to M2+ patches or M2− interpatches, in the majority of pathways (i.e. V1→LM, V1→AL, V1→RL, V1→MOs) cell bodies are not aligned to the patch/interpatch grid in L1. In contrast, M2− interpatches impose spatial order to V1→PM-, V1→RSP- and V1→LP-projecting cells.

### Somatostatin-expressing neuronal processes in V1 are localized in M2− interpatches in L1

A popular model suggests disinhibition from SST neurons underlies increased excitation of projection neurons in V1 (***Fu et al., 2014***). Prompted by our finding that M2− neurons are more robustly modulated by locomotion, we wanted to know whether axons of SST-positive neurons are denser in M2− interpatches than in M2+ patches. To find out, we immunostained tangential sections through flatmounted cortex of SST-Cre x Ai9 mice (SSTtdT) with an antibody against M2. M2 was visualized with an Alexa 647-tagged IgG and M2+ patch/M2− interpatch borders were delineated using the procedures described by ***Meier et al.*** (***2021***). Images were high-pass filtered, blurred and partitioned into six intensity quantiles of equal area. Overlays of SSTtdT and M2 showed that labeled axons in L1 were non-uniformly distributed, with a strong preference for M2− interpatches (***Figure 7a-c***). Dividing images into M2 intensity quantiles revealed that STTtdT expression is negatively correlated with M2 expression, with 4.6-times higher densities in the center of M2− interpatches than M2+ patches (***Figure 7d***; N = 5 mice; *p* < 10^-8^, r = −0.85, Pearson correlation). As a negative control, the SSTtdT images were parcellated into 5 µm-wide squares and shuffled, from which new SSTtdT intensity values were obtained for each quantile.

**Figure 7.**
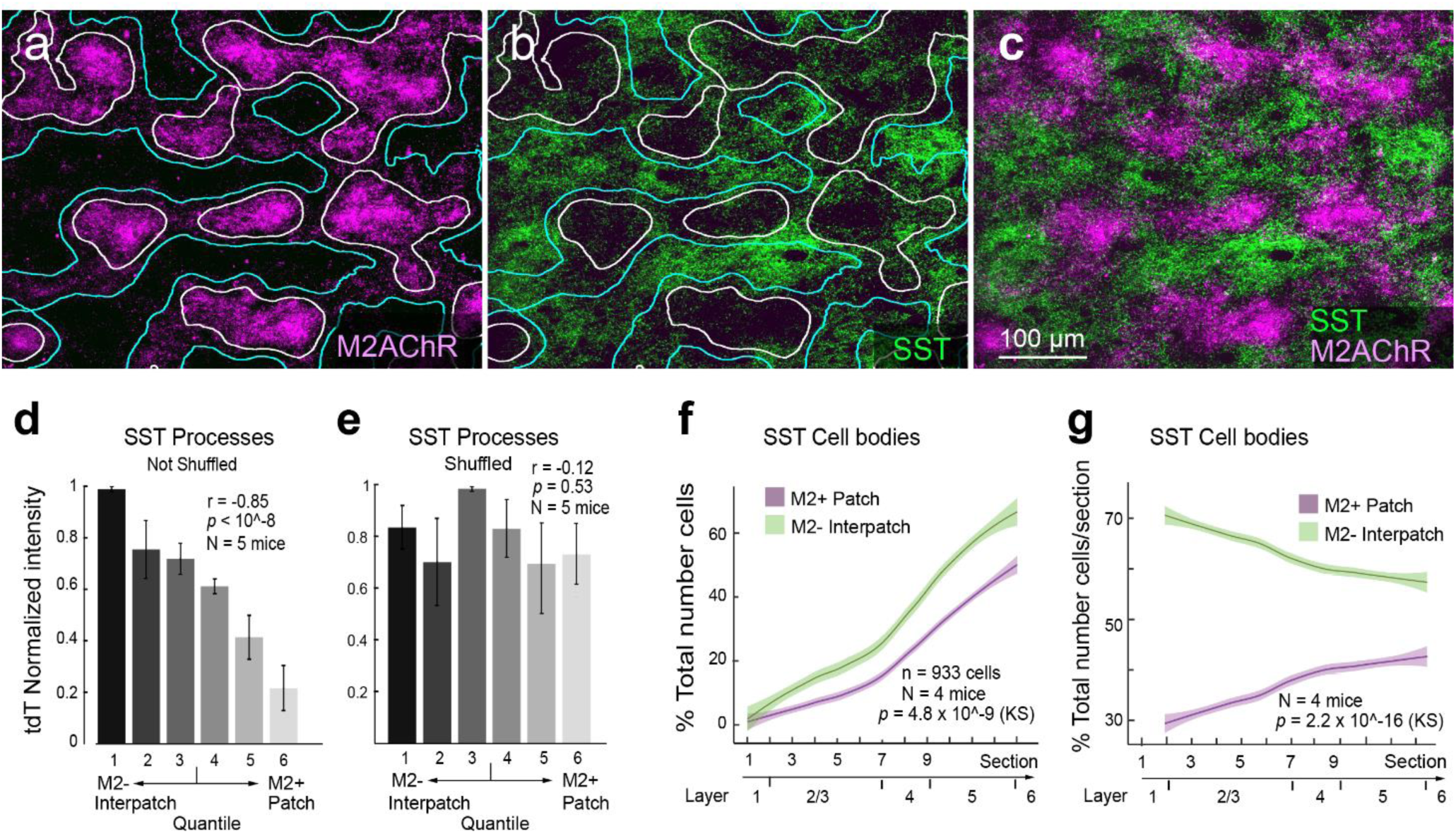
Preferential expression of SSTtdT-expressing axons and dendrites in M2− interpatches of L1 of V1 of Sst-IRES-cre x Ai9 mouse. (**a**) Tangential section through V1 shows M2+ patches (magenta, white outlines) in L1 of V1 labeled with an antibody against M2AChR. (**b**) tdT-expressing SST axons and dendrites in M2− interpatches (false colored green, blue outlines) of L1 of V1, (**c**) Overlay of panels a and b. (**d**) Intensity of SSTtdT expressing axons in different M2 quantiles, shows a strong preference for M2− interpatches. Error bars ±SEM. (**e**) SSTtdT intensity in each M2 quantile after shuffling images. (**f**) Distribution of labeled cell bodies shows that the majority of SST cells is preferentially aligned with M2− interpatches (green). (**g**) Cell counts shows that in all layers, SST cells are enriched in M2− interpatches. Shading in f and g indicates ±SEM.

The shuffled data showed no systematic relationship between SST expression and quantile (***Figure 7e***; *p* = 0.53, r = −0.12, Pearson correlation). Next, we asked whether a similar organization exists for SST cell bodies in the layers below L1. We aligned tangential sections across L2-6 to the M2 pattern in L1, and found that the percentage of the total number of SST cells was 1.5-2 times higher (n = 933 cells, N = 4 mice, *p* = 4.8 × 10^-9^, KS) underneath M2− interpatches than M2+ patches (***Figure 7f***). Similar results were found when we normalized the cell count in each module to the total number of cells in each section (***Figure 7g***). Because there are fewer SST cells in upper layers, the difference between M2− and M2+ columns was more striking in L2/3 than in the more densely populated layers 4-6 (***Wu et al., 2023***) (***Figure 7f, g***). These results show that SST cells in V1 are organized in columns preferentially aligned with M2− interpatches.

## Discussion

V1 integrates movement-related signals from multiple subcortical and cortical brain areas, which impact visual responses (***Parker et al., 2022***). Although encoding of motor variables has been found in many different cortical areas (***Musall et al., 2019; Steinmetz, et al., 2019; Stringer et al., 2019***), we discovered that locomotion-related effects on visual responses are non-uniformly distributed not only across the brain (***Chen et al., 2024***), but within a single area, V1. Our results show that locomotor effects on visual responses are more robust in M2− interpatch than M2+ patch L2/3 PCs. Consistent with this architecture, we found that apical dendrites of M2− interpatch feedforward-projecting L2/3 PCs in V1 overlap with feedback connections from LP, PM, RSP and MOs to L1, suggesting reciprocal like-to-like loops of strongly locomotion-modulated networks. The M2− associated networks interdigitate with M2+ patch-preferring loops between V1↔LM, V1↔AL and V1↔RL, in which locomotion modulation was weaker and orientation selective responses were more dominant. The modular architecture of apical dendrites of feedforward-projecting V1 neurons and overlapping feedback input from higher cortical areas suggests that the bifurcated roadmap of a dorsal and a ventral processing streams (***Wang et al., 2012***) is more complex. Our new findings suggest that the dorsal stream is divided into a medial M2− subnetwork and a lateral M2+ subnetwork, whereas the ventral stream is preferentially linked to M2+ patches (***Figure 8a***).

**Figure 8.**
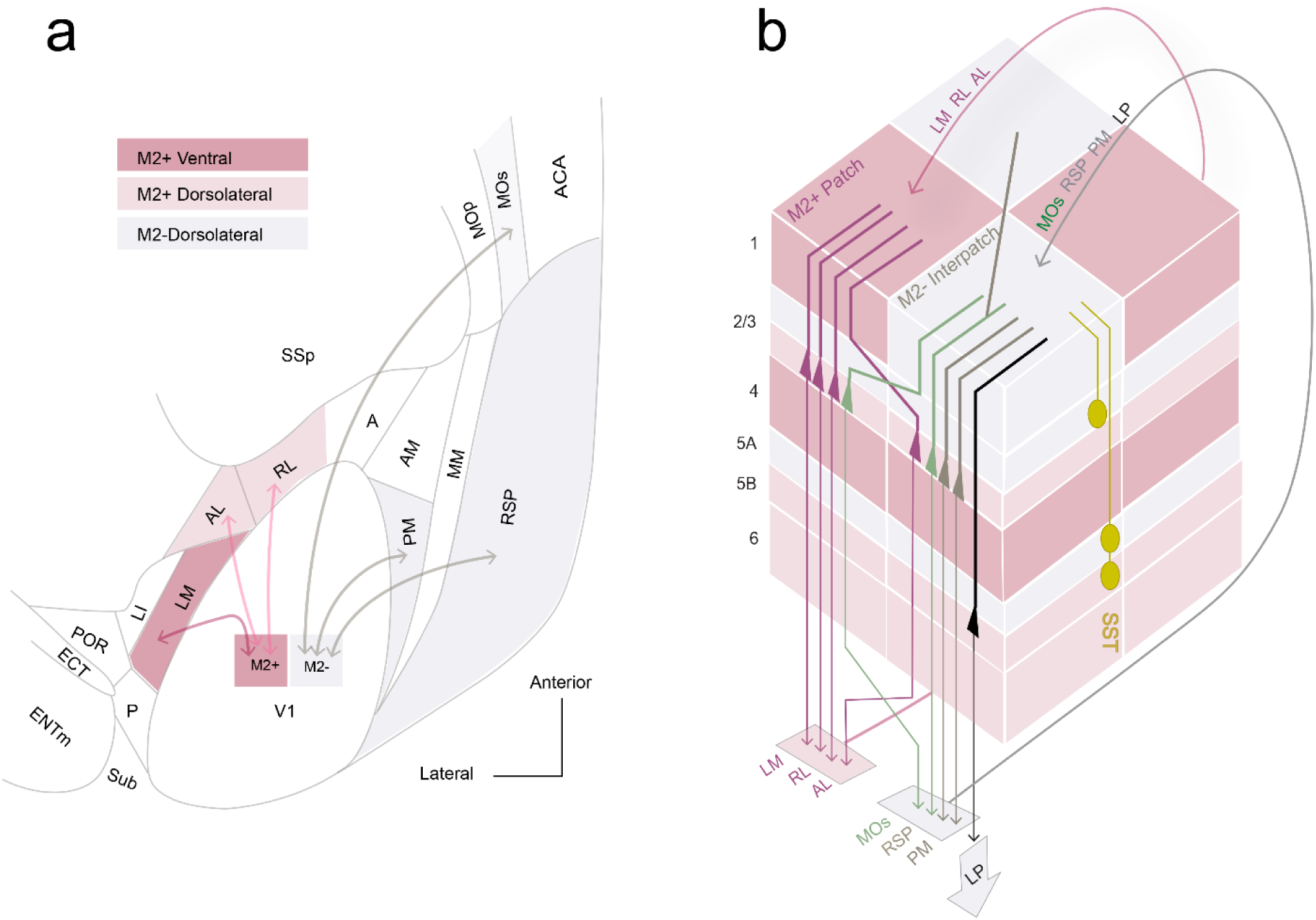
Connectivity rules of visual processing streams. (**a**) Flatmap of processing streams between V1 and higher cortical areas. The scheme is referenced to the clustering of M2+ patches (red) and M2− interpatches (grey) in L1 of V1. Each L1 module contains apical dendrites of different sets of output neurons in L2-5, and receives overlapping feedback input from corresponding higher areas to L1. The connections, represented by double arrowed lines, indicate module-selective, like-to-like reciprocal loops. M2+ patches and M2− interpatches split the dorsal and ventral intracortical processing streams into distinct sub-streams. The dorsal stream, represented by loops between V1 and AL, RL, PM, RSP and MOs, is divided into two subnetworks: the M2+ dorsolateral subnetwork (light red) composed of V1↔AL (***Ji et al., 2015; D’Souza et al., 2019***) and V1↔RL, and the M2− dorsomedial subnetwork (grey) represented by V1↔PM, V1↔RSP, and V1↔MOs. The ventral stream (dark red) is represented by the M2+ subnetwork and forms a loop between V1 and LM. For abbreviations, see Figure 4a. (**b**) Block of V1, constructed from 60-80-µm wide bundles (microcolumns) of apical dendrites (bold lines) distributed preferentially across multiple M2+ patches (red) and M2− interpatches (grey), contained within ∼300 x 300 µm slab of L1. M2 expression in layers below L1 is spatially uniform, but varies in density (grades of red shading) in layer-specific fashion. Intracortical L2/3 output PCs (triangles) project to LM, AL and RL (red) with apical dendrites (bold red lines) in M2+ patches receive overlapping feedback projections from areas LM, AL and RL, with which they form looped, like to-like connections. Output PCs (triangles) projecting to MOs (green), RSP (grey), PM (grey) and LP (black) with apical dendrites in M2− interpatches receive overlapping feedback input from areas MOs, PM, RSP and LP, with which they form looped, like to-like connections. In each pathway the dendrites are aligned with the M2+ patches or M2− interpatches in L1. In contrast, the cell bodies of L2/3 LM-, AL-, RL- and MOs-projecting neurons are randomly distributed across M2+ patches and M2− interpatches (notice red triangles aligned with M2+ and M2− domains), while the cell bodies of PM-, RSP, and LP-projecting neurons are vertically aligned with M2− interpatches (notice green triangles underneath M2+ patches and M2− interpatches). The pathway-dependent spectrum, random to clustered, of cell body distributions, sharply contrasts with the tight spatial alignment of dendrites with M2+ patches or M2− interpatches. This shows that the random distribution of LM-, AL-, RL- and MOs-projecting L2/3 neurons contrasts with the tight non-random spatial arrangement of apical dendrites in L1. The spatial organizations of L2/3 PM, RSP- and L5 LP-projecting neurons (grey, black triangles) is tighter, showing a strict alignment of apical dendrites and cells bodies with M2− interpatches. The common principle of all pathways depicted here, is that multiple types of projection neurons whose distribution may not be strictly columnar are grouped by clustered apical dendrites in M2+ patches or M2− interpatches, into distinct target-specific “output units” (***Innocenti and Vercelli, 2010*).**

### Modular architecture of locomotion modulation in V1

Movement-related signals were found in PCs of multiple sensory areas, in agreement with prior reports (***Ayaz et al., 2019; Mao et al., 2020; Salkoff et al., 2020; Yang et al., 2020***). Our recordings from *Emx1*-expressing L2/3 PCs show that locomotion-modulated cells in V1 are not randomly distributed but are aligned with M2− interpatches. M2− cell clusters are distinct from M2+ clusters, which contain a larger percentage of orientation selective cells (***Ji et al., 2015***) and show weaker locomotion modulation. The modular architecture we have discovered differs from the salt-and-pepper organization of orientation tuning observed by ***Ohki and Reid (2007)***, but is consistent with the spatial clustering of orientation selective cell bodies and dendrites reported previously (***Ji et al., 2015; Kondo et al., 2016; Maruoka et al., 2017; Ringach et al., 2016***). Our results support the findings by ***Wekselblatt and Niell (2019)*** of diverse cell populations with distinct visual and locomotion sensitivities.

Movement-related signals reach V1 through multiple pathways originating from the mesencephalic locomotor region (***Lee et al., 2014***), the dLGN (***Erisken et al, 2014***), the LP, MOs, RSP and the anterior cingulate cortex (***Bennett et al., 2019; Bouvier et al., 2020; Guitchounts et al., 2020; Harris et al., 2019; Leinweber et al., 2017***; ***Miura and Scanziani, 2022; Roth et al., 2016***). While each of these inputs project to multiple layers, locomotion-tuned boutons in L1 have only been identified in the LP pathway (***Roth et al., 2016).*** Although the tangential distribution of boutons is not known, the patchy pattern of LP projections to L1 (***D’Souza et al., 2019***) suggests that the motion-tuned boutons are enriched in M2− interpatches. In addition to locomotion, inputs from LP are also sensitive to visual motion (***Bennet et al 2019***). Such converging sensory and motor inputs may be used to compute the degree of mismatch between the speed of locomotion and optic flow, information that captures unexpected external visual motion (***Roth et al., 2016***). Cortically generated locomotion signals to L1 of V1 derive from MOs, RSP, PM and anterior cingulate cortex (***Keller et al., 2013; Polack et al., 2013***). Such signals may depolarize apical dendrites in L1 and when coincident with optic flow-related input from the LP, feedback from MOs, RSP and PM may amplify the spike output of parent L2/3 PCs (***Fisek et al., 2023; Furutachi et al., 2024; Larkum, 2007***), enabling the distinction of self-generated motion from visual motion in the environment.

### Correlated activity in cell pairs depends on module type and separation distance

To capture the functional connectivity of M2+ patch and M2− interpatch networks, we determined pairwise activity correlations between different neural populations (***Kohn et al., 2016***), and asked whether M2+ patch- and M2− interpatch-cells belong to separate subnetworks with distinct strengths of within-network correlated activities (***Cossell et al., 2015).*** We found that spontaneous activity correlations between closely spaced (< 275 µm) cells measured in the dark were greater in M2+ patch than M2− interpatch pairs (***Figure 3a***). A possible reason for this module-specific difference is the shared input from the dLGN to M2+ patches in L1 (***D’Souza et al 2019; Ji et al. 2015***). Inputs from the dLGN shell (***Meier et al., 2021***) to L1 of V1 are clustered (***D’Souza et al., 2019***) and retinotopically more precise than L1 inputs from the LP (***Roth et al., 2016***). Thus, shared dLGN inputs may overlap more tightly, and produce correlated firing in L2/3 PCs whose dendritic arbors branch in M2+ patches. However, while shared inputs can give rise to correlated activity, it should be noted that shared inputs do not universally produce the significant magnitudes of correlation which we observed (***Ecker et al., 2010***; ***Renart et al., 2010***).

During visual stimulation, noise response correlations in M2+/M2+, M2+/M2− and M2−/M2− pairs decrease over short (< 275 µm) distances of separation (***Figure 3b***). This is comparable to what has been reported in randomly recorded cell pairs of mouse V1 (***Denman and Contreras, 2014: Yu et al., 2019***), but it differs from the flat distance-dependence found in primate V1 (***Ecker et al., 2010***). Unexpectedly, we found that during locomotion, visually driven activity increases noise response correlations between cells that are > 275 µm apart and that this effect is selective for M2−/M2− interpatch pairs (***Figure 3b***). The distance-dependence of functional connectivity shows that within the core in which RFs overlap (***Ji et al., 2015***) the network is spatially uniform. However, when influences originate outside the core and provide contextual input from the RF surround (***Keller et al., 2020a***) the functional correlations are non-uniform. It is likely that these interactions are carried out in the extensive horizontal network between M2− interpatches but is absent between M2+ patches (***Burkhalter et al., 2024***).

Finding increased noise correlations between widely separated locomotion-responsive M2− interpatch cells is counter to results showing a distance-dependent decrease (***Erisken et al., 2014; Vinck et al., 2015***). One reason for this discrepancy may be that previous recordings were performed in randomly sampled pairs of neighboring cells, conditions in which firing is decorrelated despite the abundance of shared connections (***Ecker et al., 2010; Renart et al., 2010***). In contrast, the M2− interpatch cell pairs we have recorded were hundreds of microns apart. While sparse, the inputs to M2− interpatches are highly convergent (***Burkhalter et al., 2024),*** a pattern known to maximize correlated firing (***Kohn et al., 2016; Shadlen and Newsome, 1998*)**. It is more difficult to explain why during stimulus-locked motor behavior correlations between M2−interpatch (and M2+ patch) neurons separated by < 275 µm tend to be lower than in neurons which are farther apart (***Figure 3b***). One possibility is that synchronization in closely spaced neurons is low because excitation is opposed by strong synaptic inhibition (***Renart et al., 2010***). Such an arrangement may improve stimulus discriminability during spatial attention by desynchronizing activity in the RF center (***McBride et al., 2019***). Increased correlations of inputs from M2− interpatches of the RF surround during locomotion may impair encoding of visual information from select targets but increase behaviorally relevant information from the global scene (***McBride et al., 2019***).

It has been proposed that contextual influences from the RF surround are computed by a disinhibitory circuit (***Keller et al., 2020b***) which weakens surround suppression during locomotion (***Ayaz et al., 2013***). In support of this notion, we have found that SST axons (***Wu et al., 2023***) preferentially target M2− interpatches in L1 (***Figure 7a-c***), suggesting that inhibitory inputs from SST neurons (***Murayama et al., 2009; Wu et al., 2023***) overlap with apical dendrites of M2− PCs (***D’Souza et al., 2019***). Previous findings have shown that Martinotti-type SST cells are activated by locomotion (***Bugeon et al., 2022***) and are embedded in a VIP→SST→PC circuit, which strongly modulates visual responses during running (***Fu et al., 2014***). Thus, intracortical long-range input to VIP neurons (***Zhang et al., 2014***) may reduce inhibition of apical PC dendrites by SST in L1 (***Jiang et al., 2013***) and increase synchronization. It is widely held that synchronization impairs encoding of visual information (***Cohen and Maunsell, 2009; Kohn et al., 2016***). New findings, however, show that synchronous activity is more common when mice make correct behavioral decisions (***Valente et al., 2021***). Thus, learning and stimulus-locked motor activity may temporally align recurrent network dynamics in M2− interpatch subnetworks and facilitate the integration of task-relevant inputs (***Chadwick et al., 2023***).

### Modular interareal network for integrating movement-related signals in V1

Movement-related signals are known to exist in many areas of the thalamus and cortex, in which the direction of visual flow is associated with the direction of locomotion, as well as movements of the head and the eyes. Prominent examples of areas carrying motor signals are the LP, MOs and RSP, which send locomotion and head orienting movement information to V1 (***Bouvier et al., 2020; Guitchounts et al., 2020; Miura and Scanziani, 2022; Parker et al., 2022; Roth et al., 2016; Vélez-Fort et al., 2018***). Inputs are also provided by PM, which may send movement commands to V1 that are used in choice behavior (***Jin and Glickfeld, 2020***). Transforming vision into action is performed preferentially in the dorsal cortical network (***Goodale, 2011***). The enhanced drive and increased synchronization of visual responses in M2− interpatches we have found during locomotion raised the question of whether axonal projections of the dorsal stream (***Wang et al. 2012***) are preferentially targeted to M2− interpatches. More generally, could the modular architecture of L1 be a framework for segregating functionally distinct loops between feedback projections from different higher cortical areas and the LP thalamus to apical dendrites of feedforward-projecting L2/3 PVs in V1? What we found is that feedback axons overlap with either M2+ or M2− apical dendrites of feedforward-projecting neurons, showing striking like-to-like loops of connectivity. While in all loops feedback projections and apical dendrites of feedforward-projecting neurons showed preferences for specific modules, the clustering of parent cell bodies was less tight, and varied in pathway-specific fashion. For example, cell bodies in V1↔LM, V1↔RL and V1↔MOs pathways were randomly distributed, whereas in V1↔PM, V1↔RSP and V1↔LP the cells were preferentially aligned with M2− interpatches. This organization is largely consistent with the preferential locomotion sensitivity of M2− cells, except for the MOs pathway, which suggests that locomotion sensitivity dominates in a subset of M2− cells. Connections of the ventral stream area, LM, and the dorsal stream area, RL and AL (***Ji et al, 2015***) were looped through M2+ patches.

Unexpectedly, we found that the dorsal stream areas PM, RSP and MOs looped through M2− interpatches. This pattern of connectivity suggests that the dorsal network is split into a lateral M2+ network that includes LM, AL, RL, and a medial M2− subnetwork, which comprises PM, RSP and MOs (***Figure 8a***). This organization aligns with the distinct specializations of AL and RL for optic flow speed at high temporal frequency and PM for high contrast, edge-dense stimuli moving at slow speeds for drawing attention to unexpected stimuli, respectively (***Furutachi et al., 2024***; ***Han and Bonin, 2024***; ***Yu et al., 2022***). A similar branching of the dorsal network into multiple substreams for visuospatial processing has been observed in primates (***Kravitz et al., 2013***). In monkey, substreams originate from highly interconnected posterior parietal areas, which reside at the fifth level of the cortical hierarchy (***Markov et al., 2014***), and transform visuotopic information encoded in an egocentric reference frame into an object-based world-centered frame (***Snyder et al., 1998***). The corresponding hierarchy is different in mice, in which the dorsal stream branches from a single area, V1, into a lateral M2+ -associated and a medial M2− -linked substream. In mice, allocentric coordinates of moving objects may be extracted from visual flow fields ***(Sit and Goard, 2020***), and referenced to head and eye movements at a much earlier stage of the cortical hierarchy (***Bouvier et al., 2020; D’Souza et al., 2022; Guitchounts et al., 2020***). It appears that such computations are performed primarily in the M2− interpatch-linked subnetworks of V1. Responses to the direction of visual motion have been shown to be altered by the non-visual directional signals from the LP during saccades (***Miura and Scanziani, 2022***). Notably LP is connected to M2− interpatches (***D’Souza et al., 2019***) as well as to numerous higher cortical areas including the M2− targeting areas PM, RSP and MOs (***Juavinett et al., 2020***). Thus, it seems likely that similar non-visual directional feedback signals (***Bouvier et al., 2020; Guitchounts et al., 2020***) converge with visual motion direction signals in L1, and briefly alter the response preference of L2/3 M2− interpatch neurons through inputs to apical dendrites in L1. This mechanism may provide a subnetwork for factoring out external object motion during self-motion.

The low expression of the M2 receptor in M2− interpatches raises the question of whether M2− independent cholinergic mechanisms are involved in enhancing visual responses by locomotion. Because the effect is specific to M2− interpatches, other types of cholinergic receptors must be involved. One possibility are nicotinic *α*2 nAChR expressed on *Chrna2* positive Martinotti cell axons in L1 (***Hilscher et al., 2017***), a feature that coincides with the high density of SST axons in M2− interpatches. More indirect mechanisms may rely on the spatial distribution of GABAergic neurons (***Schuman et al., 2021***) in L1 M2− interpatches. For example, module-specific locomotion modulation may rely on M1 muscarinic AChR-mediated cholinergic inhibition of neurogliaform cells and their disinhibitory effect on postsynaptic PCs during prolonged activation (***Brombas et al., 2014***).

### M2+ and M2− output subnetworks

A key finding of this study is that feedback from higher cortical areas to L1 of V1 overlap with apical dendrites of L2/3 PCs, which send feedforward input to higher areas and that these reciprocal loops are selective for either M2+ patches or M2− interpatches. How can such specificity occur within the tight spacing (< 100 µm) of patches and interpatches, when individual PCs have 300 µm-wide apical tufts (***Gouwens et al., 2019; Meyer et al, 2013***)? A likely explanation is that the apical dendritic tufts are spatially non-uniform. Golgi studies have shown that dendrites ascend in < 70 µm wide bundles known as microcolumns (***Peters and Kara, 1987***). While microcolumns are consistent with an orderly vertical architecture, ***Peters and Kara (1987)*** noted that bundles were often joined by dendrites from tangentially displaced PCs, suggesting that dendrites from multiple minicolumns of cell bodies converge onto M2+ or M2− clusters. Bundled dendrites are typical features of groups of PCs which receive input from the same source and send outputs to a shared target (***Innocenti and Vercelli, 2010***). Thus, the like-to-like loops we have found suggest that M2+ patch and M2− interpatch networks each contain multiple distinct “output units” (***Figure 8b***).

## Materials and Methods

### Mice

We performed experiments using 5-10 week-old female and male Chrm2tdT-D knock-in (B6.Cg-Chrm2tm1.1Hze/J) x Emx^IRES-cre^, Chrm2tdT-D knock-in (B6.Cg-Chrm2^tm1.1Hze^/J), Sst-IRES-Cre x Ai9 B6.Cg-Gt[ROSA]26Sor^tm9(CAG-tdTomato)Hze^/J), and Ai9 B6.Cg-Gt[ROSA]26Sor^tm9(CAG-tdTomato)Hze^/J) mice. Experimental procedures were performed in accordance with the National Institutes of Health guidelines and under the approval of the Washington University Institutional Animal Care and Use Committee.

### General surgical procedures

Mice were anesthetized with intraperitoneal injections of a mixture of ketamine (86mg/kg) and xylazine (13mg/kg). Buprenorphine-SR (0.1 mg/kg, SubQ) was injected prior to surgery for analgesia. Mice were head-fixed in a stereotactic apparatus. Body temperature was monitored and maintained at 37°C. A craniotomy was made over the injection target using a dental drill. Viral injections were delivered via glass micropipettes (tip diameter 20µm) attached to a Nanoject II pump. For all cortical injections, two injections were made, at 0.3mm and 0.5mm below the pial surface. Pipettes were kept in place for 5 min after each injection to allow for diffusion of the tracer. In cases which did not receive a head plate implant, the scalp was stapled and secured with wound clips.

### Surgery for GCaMP imaging

Emx^IRES-cre^ mice were crossed with Chrm2tdT-D knock-in (BG6.Cg-Chrm2^tm1.1Hze^)/J (Chrm2tdT) mice. Expression of Cre in Emx1-expressing cells allowed for targeting of AAV.hSyn.Flex.GCaMP6f to PCs. Chrm2tdT labeled M2 receptors, enabling visualization of M2+ patches and M2− interpatches. Under anesthesia, the scalp was retracted, and transcranial imaging of Chmr2tdT expression was performed with a stereomicroscope (Leica MZ16F) equipped with fluorescence optics (excitation 560 nm, emission 620 nm) to determine the location of V1. V1 was visible through the skull as a triangular region of high Chmr2tdT expression (***Meier et al., 2021***). Fluorescence images and brightfield images of blood vessels were acquired and overlaid to guide subsequent surgical procedures. A 3mm-diameter craniotomy was made over the center of V1 in the left hemisphere using a dental drill. This involved thinning the skull around the borders of the 3mm target area, then removing a circular flap of skull, exposing the surface of the brain. Four injections (46nl each, 0.3-0.5mm below the surface) were performed at different coordinates within V1 (in mm anterior to transverse sinus/lateral to midline: 1.0-1.9/2.4-3.3). Transparent silicone adhesive (Kwik-Sil) was applied to the surface of the exposed brain. A 3mm-diameter circular cover slip was placed on top of the Kwik-Sil, with the edges of the cover slip contacting the skull at the perimeter of the craniotomy. A layer of dental cement was then applied to the rim of the cover slip and surrounding bone, rigidly securing the cover slip to the skull. An aluminum head plate containing attachments for head fixation during live imaging was affixed to the skull with dental cement.

### 2-photon imaging

After surgery, 3 weeks were allowed to elapse for AAV-mediated gene transduction. Mice were then habituated to the recording apparatus by head-fixing them on a cylindrical running wheel (***Figure 1b***). Two 30-minute head-fixed habituation sessions were performed in the two days before recording. During each recording session, GCaMP6f signals from L2/3 Emx1-expressing neurons were acquired. An Ultima 2-photon recording system (Prairie Technologies) and Olympus BX61W1 microscope were used with a 20x objective (Olympus UMPlanFL, NA 1.0). A mode-locked laser (Mai Tai DeepSee Ti: Sapphire, Spectra-Physics) was used for 920um excitation and GCaMP6f signals were acquired through a green filter (525 nm center wave length/ 70nm band width). One plane was acquired each session at a sample rate of 3Hz, with a field of view of 0.5 x 0.5mm. The velocity of the running wheel was recorded during the entire recording session and digitized at 10kHz, to determine the onset/offset of locomotion and free-running speed of the mouse during calcium recordings.

Analysis of GCaMP6f recordings of neuronal activity were performed using Suite2p analysis software (***Pachitariu et al., 2017***) and custom programs in MATLAB, adapted from ***Arriaga and Han (2017)***. First, cross-correlation registration and rigid transformation of images in the time-series was performed to eliminate movement artifacts in the XY plane (***Dombeck et al., 2010; Arriaga and Han, 2019***). Candidate cells were then automatically extracted based on local temporal pixel intensity correlations. For analysis, cells were manually selected based on maximum and mean fluorescence. Manual selection included rejection of out-of-focus cells and axons and dendrites separated from somas. For each selected cell, fluorescence intensity of the neuropil in a 20 µm annulus surrounding the cell body was subtracted from the fluorescence intensity of the cell to determine a value for F. ΔF/F was then calculated with a 30-second sliding time window, using the 8^th^ percentile of fluorescence intensity (***Williams et al., 2019***).

### Visual stimuli

Stimuli were presented on a γ-corrected computer monitor (AOC 27B1H, 34 x 61 cm, 60 Hz refresh rate, 32.1 cd/m^2^ mean luminance) which was placed 33 cm away from the right eye at a 45° angle to the body axis, subtending 54.5° x 85.5° (vertical x horizontal) of visual space. The screen center was elevated to be level with the eye. The Psychophysics Toolbox for MATLAB was used for the creation and presentation of stimuli (***Brainard, 1997***). All stimuli consisted of circular square-wave drifting gratings at 100% contrast which were warped to simulate presentation on a sphere with its center at the mouse’s eye (***Marshel et al., 2011***). At the beginning of each session, RF mapping was performed by sequentially presenting a 20° diameter square wave grating at randomly selected locations in a 6 column by 4 row grid. The grid was centered on the visual field of the right eye and grid locations were spaced 15° apart center-to-center. For each recorded cell, responses to each trial were computed as the mean ΔF/F during the 4 seconds of stimulus presentation. A 2-dimensional Gaussian curve was then fit to the response profile of each cell (using the MATLAB ‘polyfit’ function), which was used to compute the center of its receptive field. The average X and Y coordinates of RFs of all responsive cells was computed, and this location was used for centering all stimuli in the remainder of the recording session. The stimulus screen was then turned off, and 10 minutes of spontaneous activity was recorded.

Next, the spatiotemporal tuning of cells was determined by presenting 100% contrast square wave gratings with a range of spatial frequencies (SF), temporal frequencies (TF), and orientations (OS). Peak luminance of light stripes was 62.2 cd/m^2^ and minimum luminance of dark stripes was 2.3 cd/m^2^. A fixed location for these stimuli was chosen to match the RFs computed in the first stimulus block. Eight SFs ranging from 0.01-1.6 c/deg, 7 TFs ranging from 0.1-13 Hz, and 7 equally spaced orientations (51° increments) were presented, with 10 repetitions per unique stimulus. When SF and TF were not being tested, they were fixed at 0.03 cycles per degree and 2Hz (drift speed = 67°/sec). To determine the preferred SF and TF and OS of each cell, a log Gaussian function was fit to SF and TF responses and a von Mises function was fit to OS responses (***Gao et al., 2010***). When comparing SF, TF, orientation, and locomotion tuning across the population (***Figure 2, Figure 2 Supplement 2.1, and Figure 2 Supplement 2.2***), we excluded from the analysis all cells which had RFs that were not covered by the stimuli used to test for SF, TF, and orientation tuning during the third phase of the experiment. When analyzing SF, TF, and orientation HWHM and peak (***Figure 2 Supplement 2.2***), cells were excluded if they did not show significant (α = 0.05, ANOVA) tuning for the stimulus parameter (SF, TF, or orientation) in question, as these non-tuned cells may have spurious extreme values for HWHM and peak. For a minority of cells (approximately 10% for each of SF/TF/orientation), HWHM could not be accurately estimated using our fitting functions within the range of parameters values that had been presented, and these cells were excluded from HWHM analyses. When comparing SF, TF, and orientation HWHM during rest vs. locomotion (***Figure 2 Supplement 2.2 a-c***), cells were also excluded if the session in which they were recorded did not have at least 3 rest trials and 3 locomotion trial during which the stimulus parameter in question was being varied.

A locomotion modulation index (LMI) was computed to determine each cell’s response gain due to locomotion. To compute the LMI, trials were first classified as locomotion trials if mean forward locomotion during stimulus presentation was > 1 cm/sec, or stationary trials if mean forward locomotion was < 1 cm/sec (***Pakan et al., 2016***). The locomotion modulation index was then computed as (R_L_ – R_S_) / (R_L_ + R_S_), where Response during Locomotion R_L_ = mean ΔF/F on locomotion trials, Response while Stationary R_S_ = mean ΔF/F on stationary trials.

### Aligning recorded cells with M2+ patches and M2− interpatches

After completing four to five imaging sessions (1 session/day, using non-overlapping regions of V1), mice were perfused through the heart with 1% paraformaldehyde (PFA). Cortex was flatmounted, postfixed in 4% PFA, cryoprotected in 30% sucrose, and sectioned horizontally at 40 µm on a freezing microtome. Images of the cortical window that were previously acquired *in vivo* were used to locate the recorded regions in *ex vivo* sections (***Figure 1a***). Sections were imaged at 2-20 x magnification with an epifluorescence microscope (Nikon 80i) equipped with GFP (excitation 490 nm, emission 510-538 nm) and tdTomato (excitation 550 nm, emission 570-720 nm) optics, to determine the M2+ patch pattern of recorded regions. To determine M2+ patches from Chmr2tdT fluorescence, images were first spatially normalized by dividing the intensity of each pixel by the average intensity from a circle with 100 µm radius surrounding it (***D’Souza et al., 2019; Meier et al., 2021***). Images were then blurred with a circular averaging filter of 30 µm radius. The image was then divided into six quantiles based on the resulting pixel intensities, with the top two quantiles considered to be M2+ patches and the bottom two considered to be M2− interpatches. Automated determination of quantile boundaries was determined with custom MATLAB scripts.

Landmarks from the *in vivo* recording sessions, including blood vessels and GFP fluorescence, were identified in the *ex vivo* section (***Figure 1c-e***). The *ex vivo* and *in vivo* images were aligned based on these landmarks. To account for tissue distortion occurring between recording and sectioning, fiducial points were assigned in *in vivo* and *ex vivo* images, which were used for alignment. Warping was performed using a projective transformation via the MATLAB ‘fitgeotrans’ function. Recorded cells were assigned as M2+ patch or M2− interpatch cells depending on which quantile they aligned with (***Figure 1f-h***).

### Correlation Analysis

Activity correlations in darkness (***Figure 3a***) were computed by finding the Pearson correlation coefficient between ΔF/F of each pair of neurons within a session during the 10 minute dark period. To compute stimulus response noise correlations (***Figure 3b-d***), a single response value was first computed for each trial for each cell by taking the average ΔF/F of that cell during the 4 seconds of stimulus exposure. For each unique stimulus condition *φ* (identical SF, TF, and OS), the Pearson correlation coefficient between the responses of each of a pair of cells across the 10 repetitions of the unique stimulus was computed (***Ecker et al., 2010; Montijn et al., 2014***). The mean correlation coefficient over all unique stimulus parameters (22 unique stimuli from 8 SFs, 7 TFs, and 7 OS) was then used as the overall response noise correlation for the cell pair. Thus, each cell pair *i, j* was assigned a noise correlation value *ρ_i, j_*

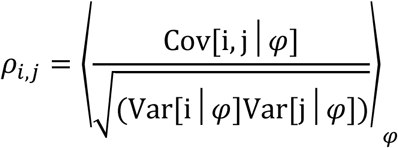

where i and j are the vectors of response magnitudes from cells *i, j* to each unique stimulus *φ*.

To determine whether cells were locomotion-responsive, the Pearson correlation between the ΔF/F trace of the cell and the locomotion velocity during the 10 minute dark period was computed. Cells were considered significantly locomotion responsive (***Figure 3c, d***) if the *p*-value of this correlation was < 0.05.

### Immunostaining

To visualize expression of M2 in Ai9 and SST-Cre x Ai9 mice, immunostaining was performed on sections cut on a freezing microtome at 40µm using an antibody against M2 muscarinic acetylcholine receptor. First, sections were preincubated in a blocking solution (0.1% TritonX-100, 10% normal goat serum, PB). Next, incubation was performed in primary rat anti-M2 antibody (1:500, MAB367 [Millipore], 48h at 4°C) and reacted with Alexa-647-labeled goat anti-rat secondary antibody (1:500, A21247, Invitrogen). Epifluorescence microscopy was then used to image the pattern of M2 expression in different layers.

### Anterograde and retrograde viral tracing

We examined the distribution of axons, cell bodies and dendrites projecting to and from M2+ patches and M2− interpatches of V1 by anterograde and retrograde viral tracing in Ai9 mice. Inputs and outputs of different structures were labeled by injecting a cocktail of AAV2/1-hSyn-EGFP.WPRE.bGHe (46nl, University of Pennsylvania, Vector Core) and rAAV2.Retro-CAG.Cre (46nl, University of North Carolina, Vector Core) into (in mm, anterior-posterior/lateral/deep): dLGN (−2.35 post bregma/2.15/2.7 deep), LP (−1.85 post bregma/1.25/2.75), LM (1.4 anterior to transverse sinus (ts)/4.0/0.3 and 0.5), RL (2.3 ant ts/3.2/0.3 and 0.5), PM (1.9 ant ts/1.6/ 0.3 and 0.5), RSP (−2.75 post bregma/1.2/0.3 and 0.5), and MOs (−0.15 post bregma/0.5/0.3 and 0.5). Three weeks later, mice were perfused with 1% PFA, the cortex was flatmounted, postfixed in 4% PFA, cryoprotected in 30% sucrose, and cut on a freezing microtome at 40 µm in the tangential plane. Immunostaining against M2 was performed in tangential sections to delineate M2+ patches, M2− interpatches and areal borders (***Gămănuţ et al., 2018***). Sections were wet-mounted on slides and images were acquired under a fluorescence microscope at 2-40 X magnification, which were used to delineate M2+ patch borders and identify labeled cell bodies axons and dendrites. The remainder of the brain was then separated from the cortex. In cases with dLGN or LP injections, the remainder of the brain was sectioned at 40 µm in the coronal plane, and fluorescence images were obtained to assess the location of the subcortical injection site.

### Quantification of axons, dendrites and cell bodies in M2+ patches and M2− interpatches

The patch/interpatch pattern in L1 of V1 was visualized by M2 immunostaining. Fluorescence images of tangential sections through L1 were high-pass filtered and blurred, and labeling was divided into 6 equally sized intensity quantiles (***D’Souza et al., 2019***; ***Sincich and Horton, 2002; Sincich and Horton, 2005***). The top three quantiles were designated as M2+ patches while the bottom three quantiles were denoted as M2− interpatches. Anterogradely labeled axons of inputs from LP, LM, RL, PM, RSP and MO to L1, and retrogradely labeled apical dendritic tufts of LP-, LM-, RL-, PM-, RSP- and MOs-projecting cells in L1 were identified using morphological criteria (***Karimi et al., 2020***). To determine the strength of axons and dendrites in M2+ patches and M2− interpatches, we used a custom MATLAB script to measure the pixel values of fluorescence intensity in ROIs (550 um x 400 um) and subtracted the unlabeled background. The mean and standard error of fluorescence intensity within each M2+ and M2− quantile was determined across 3 ROIs per section, averaged across mice and plotted as histograms. The mean intensities of axonal and dendritic labeling in M2+ patches and M2− interpatches were compared across quantiles using a Pearson correlation.

For counting retrogradely labeled cell bodies aligned with M2+ patches and M2− interpatches in different layers, we first plotted contours outlining M2+ patches (top 3 quantiles) and M2−interpatches (bottom 3 quantiles) in L1. We then aligned sections L2-6 using blood vessels as landmarks and transferred the patch and interpatch contours from L1. Retrogradely labeled cells were assigned to M2+ patches and M2− interpatches, disregarding cells intersected by contour lines, using the multipoint selection tool of ImageJ (NIH). The mean and standard error of the number of cells was computed across 3 ROIs per section, averaged across mice and plotted as smooth curves. The Kolmogorov-Smirnov (KS) test was used to compare the total number of cells across layers. Tangential section number was converted into depth to extract layers identified in coronal sections (***Paxinos and Franklin, 2013***). This translation of tangential section number into layers was validated recently from the differential laminar expression patterns of parvalbumin in L2-4 and Ctip 2 (COUP-TF-interacting protein 2) in L5-6 (***Burkhalter et al., 2024***).

### Spatial distribution of SST-expressing cells and processes

To determine the strength of SST expression in M2 intensity quantiles, M2 expression images were high pass filtered and blurred. M2 images were then divided into 6 intensity quantiles. The average optical density of axons in ROIs (µm x µm, 10X, 3 per section) within each quantile were then found. The mean and standard error of intensity within each quantile was computed across multiple ROIs per section and subjects and plotted as histograms. Pearson correlation coefficient and *p*-value between quantile and intensity was computed, to determine whether axonal SST expression were associated with M2+ patches or M2− interpatches. SSTtdT expression was also compared against a shuffled distribution. SSTtdT images were downsampled by averaging all pixel intensities within a 5 x 5 µm square, the locations of these units were shuffled, then mean intensity within each M2 quantile was measured, using the original M2 quantile borders. Analysis of the correlation between quantile and shuffled intensity was performed, as with the non-shuffled images. SST cells bodies were counted using the procedures described for projection neurons.

## Supplemental Figures

**Figure 1 Supplement 1.**
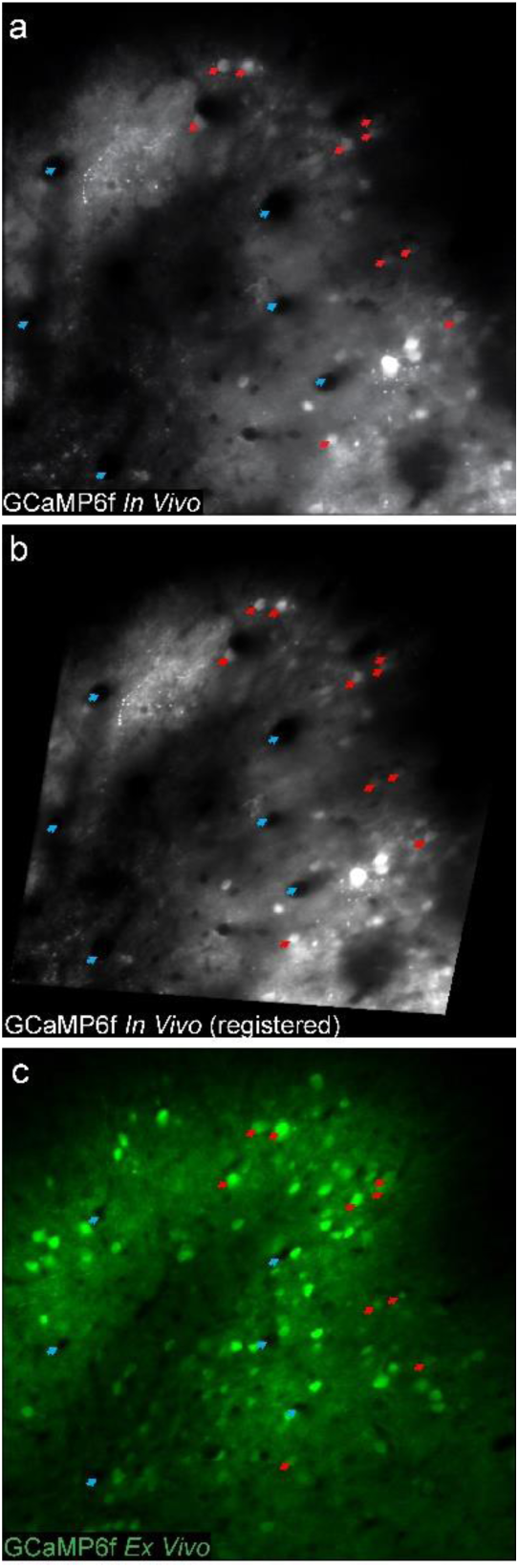
Demonstration of image registration of *in vivo* recording planes with *ex vivo* sections. (**a**) Non-registered image of time-averaged GCaMP6f fluorescence across a session in the recording plane. Cyan arrows point to blood vessels and red arrows point to GCaMP6f-labeled cells which were clearly identifiable in both this *in vivo* image and the corresponding *ex vivo* section (panel **c**). These points were manually chosen in both images and used as fiducial points for registration. (**b**) Registered version of the *in vivo* image. Registration consisted almost entirely of rotation, with very little distortion or change in relative distances between points in the *in vivo* image. The Matlab function ‘fitgeotrans’ was used by inputting point pairs, including those labeled with red arrows, from the non-registered *in vivo* image (panel **a**) and the *ex vivo* image (panel **c**). *In vivo* landmarks were used as “moving points” and *ex vivo* landmarks were used as “fixed points.” (**c**) GCaMP6f fluorescence in *ex vivo* section at same location as *in vivo* recording. Cyan and red arrows are in the same locations and point to the same anatomical structures in this section and in the registered *in vivo* image (panel **b**), demonstrating that the registration process was successful in precisely aligning locations across the two conditions.

**Figure 2 Supplement 2.1.**
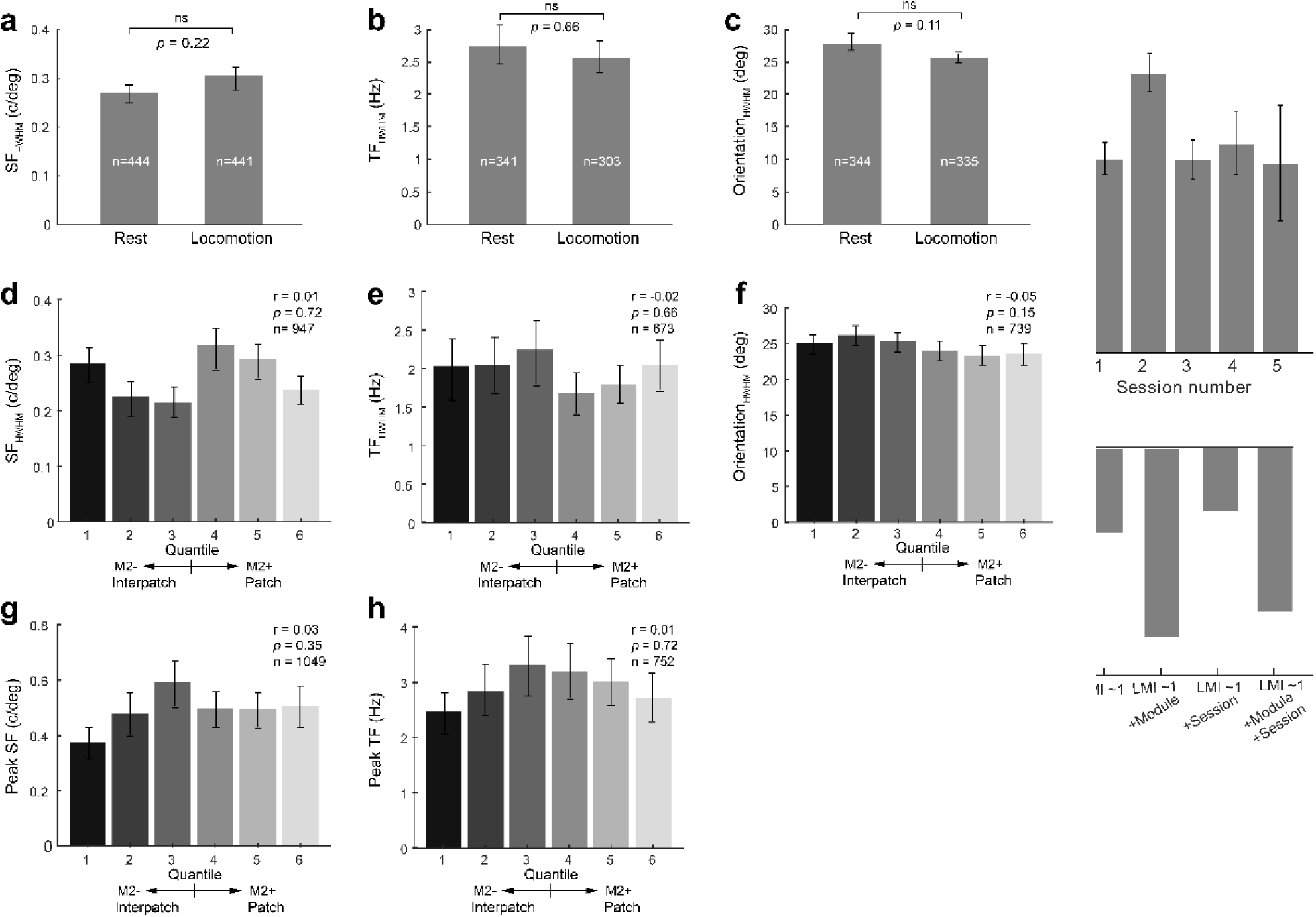
Effect of recording session number on locomotion modulation. (**a**) Mean LMI across cells in all mice according to recording session number. Cells were divided into 5 groups depending on how many experimental sessions the mouse had experienced prior to that cell being recorded. Experience in prior recording sessions neither systematically increased nor decreased LMI of cells (r = 0.006, *p* = 0.882, Pearson correlation). (**b**) Relative goodness of fit of linear models for LMI which either included or excluded session number as an independent variable. Akaike Information Criterion (AIC) was used to compare model suitability. Using session number as the only predictor (3^rd^ model) was inferior to a model using only a fixed intercept (1^st^ model). Similarly, adding session number as a predictor (4^th^ model) to a model using M2 module as a predictor (2^nd^ model) reduced the model suitability.

**Figure 2 Supplement 2.2** Effects of locomotion and M2 module on visual tuning parameters of neural responses. (**a-c**) Half-width at half maximum (HWHM) of spatial frequency (SF), temporal frequency (TF), or orientation tuning computed from locomotion (mean forward velocity > 0.1cm/s) and stationary trials (mean forward velocity < 0.1cm/s). For each cell, a tuning curve while stationary or during running were computed. Locomotion state did not change the HWHM of any stimulus parameter (*p* > 0.05, paired t-test). (**d-f**) HWHM of SF, TF and orientation tuning of cells in M2+ patches and M2− interpatches. M2+ patch and M2− interpatch cells did not show differences in HWHM for any tuning parameter *(p* > 0.05, t-test). (**g-h**) Peak SF and TF of cells in M2+ patches and M2− interpatches. M2+ patch and M2− interpatch cells did not show differences in peak SF or TF (*p* > 0.05, Pearson correlation).

**Figure 5 Supplement 1.**
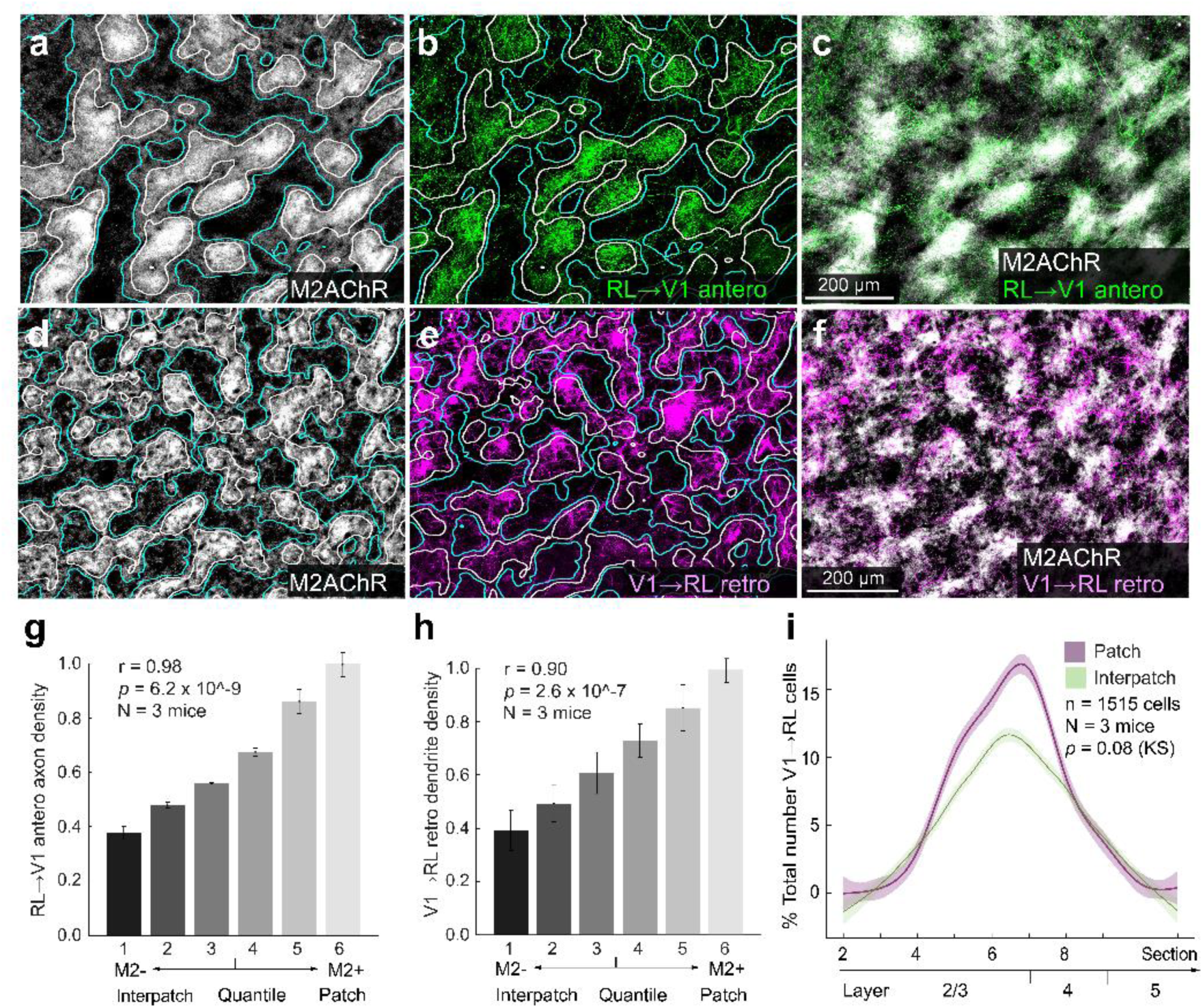
Modular organization of connections between V1 and RL. (**a**) Tangential section through L1 of V1 of Ai9 mouse stained with antibody against M2. M2+ patches (white) are outlined by white contours, M2− interpatches by cyan lines. (**b**). Anterogradely AAV-labeled RL→V1 axons (green) in L1 of V1 shows preferential termination in M2+ patches. (**c**) Overlay of panel a and b. (**d-f**) Apical dendrites of retrogradely AAV-labeled V1→RL-projecting neurons in L1 preferentially branch in M2+ patches. (**g**) Labeling density of axons in different M2 quantiles. Pearson correlation (r), error bars ±SEM. (**h**) Labeling density of dendrites in different M2 quantiles. Pearson correlation (r), error bars ±SEM. (**i**) Percent of retrogradely AAV-labeled V1→RL-projection neurons in M2+ patches (magenta) and M2− interpatches (green) in different layers of V1. Shading ±SEM. KS test.

**Figure 6 Supplement 1.**
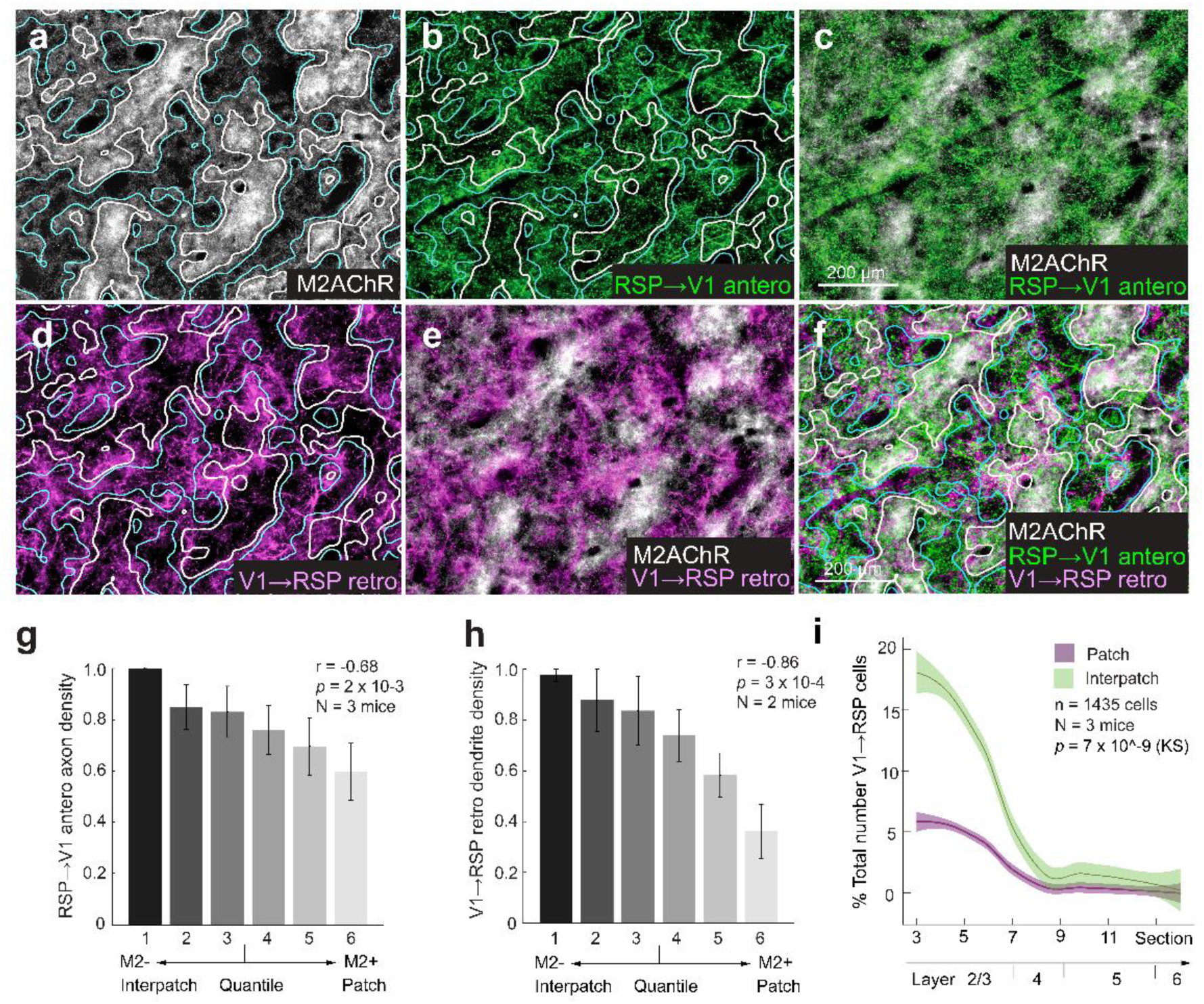
Modular organization of connections between V1 and RSP. (**a**) Tangential section through L1 of V1 of Ai9 mouse stained with antibody against M2. M2+ patches (white) are outlined by white contours, M2− interpatches by cyan lines. (**b**, **c**). Anterogradely AAV-labeled RSP→V1 axons (green) in L1 of V1 shows preferential termination in M2− interpatches. (**d-f)** Apical dendrites of retrogradely AAV-labeled V1→RSP-projecting neurons in L1 preferentially branch in M2− interpatches. (**g**) Labeling density of axons in different M2 quantiles. Pearson correlation (r), error bars ±SEM. (**h**) Labeling density of dendrites in different M2 quantiles. Pearson correlation (r), error bars ±SEM. (**i**) Percent of retrogradely AAV-labeled V1→RSP-projection neurons in M2+ patches (magenta) and M2− interpatches (green) in different layers of V1. Shading ±SEM. KS test.

**Figure 6 Supplement 2.**
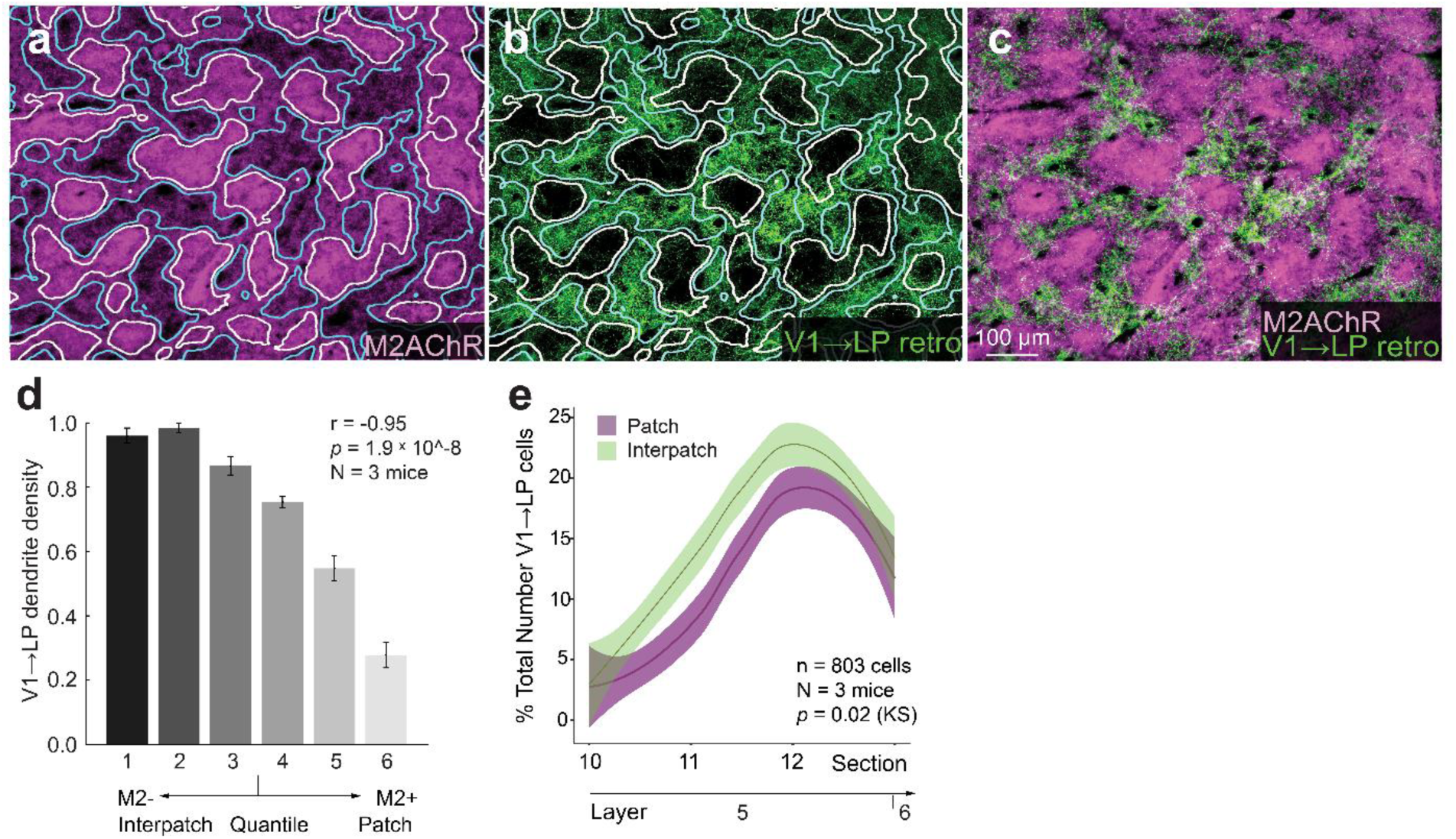
Modular organization of connections between LP and V1. (**a**) Tangential section through L1 of V1 of Ai9 mouse stained with antibody against M2. M2+ patches (magenta) are outlined by white contours, M2− interpatches by cyan lines. (**b**, **c)** Apical dendrites (green) of retrogradely AAV-labeled V1→LP-projecting L5 neurons preferentially branch in M2− interpatches of L1. (**d**) Labeling density of dendrites in different M2 quantiles shows that dendritic branches are denser in M2− interpatches. Pearson correlation (r), error bars ±SEM. (**e**) Retrogradely labeled V1→LP-projecting cell bodies show that cell bodies in L5 are preferentially (*p* = 0.02, KS) aligned with M2− interpatches (green). Shading ±SEM. KS test.

## Acknowledgements

We would like to thank James Fitzpatrick for advice in microscopy and image analysis.

## Additional Information

## Author contributions

Author contributions: AM and AB designed research. AM collected and analyzed physiological data. The anatomical data were collected and analyzed by AM, AB, PB, RD and WJ. EH provided advice in 2P imaging and valuable comments on the manuscript. AM and AB wrote the paper.

## Autor

Ethics and Animal experimentation: All experimental procedures were approved by the Animal Care and Use Committee at the Washington University (protocol numbers 20130104, 20190094, 22-0102) and conformed to the guidelines set by the National Institutes of Health.

## REFERENCES

Adesnik H, Burns W, Taniguchi H, Huang ZJ, Scanziani M. 2012. A neural circuit for spatial summation in visual cortex. Nature 490: 226–231. DOI: 10.1038/nature11526, PMID:23060193

Arriaga M, Han, EB. 2017. Dedicated hippocampal inhibitory networks for locomotion and immobility. J Neurosci 37: 9222–9238. DOI: 10.15523/JNEUROSCI.1076-17.2017, PMID:28842418

Arriaga M, Han EB. 2019. Structured inhibitory activity dynamics in new virtual environments. eLife 8: e47611. DOI: 10.7554/eLife.47611, PMID: 31591960.

Ayaz A, Saleem AB, Schölvinck ML, Carandini M. 2013. Locomotion controls spatial integration in mouse visual cortex. Curr Biol 23: 890–894. DOI: 10.1016/j.cub.2013.04.012, PMID: 23664971

Ayaz A, Stäuble A, Hamada M, Wulf MA, Saleem AB, Helmchen F. 2019. Layer-specific integration of locomotion and sensory information in mouse barrel cortex. Nat Common 10: 2585. DOI: 10.1038/s41467-019-10564-8, PMID: 31197148

Bennett C, Arroyo S, Hestrin S. 2013. Subthreshold Mechanisms Underlying State-Dependent Modulation of Visual Responses. Neuron 80: 350–357. DOI : 10.1016/j.neuron.2013.08.007, PMID: 24139040

Bennett C, Gale SD, Garrett ME, Newton ML, Callaway ED, Murphy GJ, Olsen SR. 2019. Higher-order thalamic circuits channel parallel streams of visual information in mice. Neuron 102: 477–492. DOI: 10.1016/j.neuron.2019.02.010, PMID 30850257

Bermejo F, Hug MX, Di Paolo EA. 2020. Rediscovering Richard Held: Activity and passivity in perceptual learning. Front Psychol 11: 844. DOI: 10.3389/fpsyg. 2020.00844. eCollection 2020, PMID: 32508708

Bouvier G, Senzai Y, Scanziani M. 2020. Head movements control the activity of primary visual cortex in a luminance-dependent manner. Neuron 108: 500–511. DOI :10.1016/j.neuron.2020.07.004, PMID : 32783882

Brainard DH. 1997. The Psychophysics Toolbox. Spat Vis 10: 433–436, PMID: 917695

Brombas A, Fletcher LN, Williams SR. 2014. Activity-dependent modulation of layer 1 inhibitory neocortical circuits by acetylcholine. J Neurosci 34: 1932–1941. DOI: 10.1523/JNEUROSCI.4470-13.2014, PMID: 24478372

Bugeon S, Duffield J, Dipoppa M, Ritoux A, Prankerd I, Nicoloutsopoulos D, Orme D, Shinn A, Peng H, Forrest H, Viduolyte A, Reddy CB, Isogai Y, Carandini M, Harris KD. 2022. A transcriptomic axis predicts state modulation of cortical interneurons. Nature 607: 330–338. 10.1038/s41586-022-04915-7, PMID: 35794483

Burkhalter A, Ji W, Meier AM, D’Souza RD. (2024). Modular horizontal network within mouse primary visual cortex. Front Neuroanat 18: 1364675. DOI: 10.3389/fana.2024.1364675, PMID: 38650594

Chadwick A, Kahn AG, Poort J, Blot A, Hofer SB, Mrsic-Flogel TD, Sahani M. 2023. Learning shapes cortical dynamics to enhance integration of relevant sensory input. Neuron 111: 106–120. DOI : 10.1016/j.neuron.2022.10.001, PMID: 36283408

Carvalho MM, Tanke N, Kropff E, Witter MP, Moser M-B, Moser EI. 2020. A brainstem locomotor circuit drives the activity of speed cells in the medial entorhinal cortex. Cell Reports 32:108123. DOI: 10.1016/celrep.2020.108123, PMID: 32905779

Chen G, King JA, Burgess N, O’Keefe J. 2013. How vision and movement combine in the hippocampal place code. Proc Nat Acad Sci USA 110:378–383. DOI: 10.1073/pnas.125834110, PMID: 23256159.

Chen S, Liu, Y, Wang, ZA, Colonell, J, Liu DL, Hou H, Tien N-W, Wang T, Harris T, Druckmann S, Li N, Svoboda K. 2024. Brain-wide neural activity underlying memory-guided movement. Cell 187: 676–691. DOI: 10.1016/j.cell.2023.12.035, PMID: 38306983

Cohen MR, Maunsell JHR. 2009. Attention improves performance primarily by reducing interneuronal correlation. Nat Neurosci 12: 1594–1600. DOI: 10.1038/nn.2439, PMID: 19915566

Cossell L, Iacaruso MF, Muir DR, Houlton R, Sader EN, Ko H, Hofer SB. Mrsic-Fogel TD. 2015. Functional organization of excitatory synaptic strength in primary visual cortex. Nature 518: 399–403. DOI: 10.1038/nature14182, PMID: 25652823

Dadarlat MC, Stryker MP. 2017. Locomotion enhances neural encoding of visual stimuli in mouse V1. J Neurosci 37: 3764–3775. DOI: 10.1523/JNEUROSCI.2728-16.2017, PMID: 28264980

Denman DJ, Contreras D. 2014. The structure of pairwise correlation in mouse primary visual cortex reveals functional organization in the absence of an orientation map. Cereb Cortex 24: 2707–2720. DOI: 10.1093/cercor/bht128, PMID: 23689635

Dipoppa M, Ranson A, Krumin M, Pachitariu M, Carandini M, Harris KD. 2018. Vision and locomotion shape the interactions between neuron types in mouse visual cortex. Neuron 98, 602–615.e8. DOI: 10.1016/j.neuron.2018.03.037, PMID : 29656873

Dombeck DA, Harvey CD, Tian L, Looger LL, Tank DW. 2010. Functional imaging of hippocampal place cells at cellular resolution during virtual navigation. Nat Neurosci 13: 1433–1440. 10.1038/nn.2624, PMID: 20890294

D’Souza RD, Bista P, Meier AM, Ji W, Burkhalter A, 2019. Spatial Clustering of Inhibition in Mouse Primary Visual Cortex. Neuron 104: 588–600. DOI://https://10.1016/j.neuron.2019.09.020, PMID: 31623918

D’Souza RD, Wang Q, Ji W, Meier AM, Kennedy H, Knoblauch K, Burkhalter A. 2022. Hierarchical and nonhierarchical features of the mouse visual cortical network. Nat Commun 13: 503. DOI: 10.1038/s41467-022-28035-y, PMID: 35082302

Ecker AS, Berens P, Keliris GA, Bethge M., Logothetis NK, Tolias AS. 2010. Decorrelated neuronal firing in cortical microcircuits. Science 327:584–587. DOI: 10.1126/science.1179867, PMID: 20110506

Erisken S, Vaiceliunaite A, Jurjut O, Fioini M, Katzner S, Busse L. 2014. Effects of locomotion extend throughout the mouse early visual System. Curr Biol 24: 2899–2907. DOI: 10.1016/j.cub.2014.10.045, PMID: 25484299

Fajen BR, Matthis JS. Visual and non-visual contributions to the perception of object motion during self-motion. 2013. PLOS One 8: e55446. DOI: 10.1371/journal.pone.0055446, PMID : 23408983

Fisek M, Herrmann D, Egea-Weiss A, Cloves M, Bauer L, Lee T-Y, Russell LE, Häusser M. 2023. Cortico-cortical feedback engages active dendrites in visual cortex. Nature 617:769–776. DOI: 10.1038/s41586-023-06241-y, PMID: 37280368

Fu Y, Tucciarone JM, Espinosa JS, Sheng N, Darcy DP, Nicoll RA, Huang ZJ, Stryker MP. 2014. A cortical circuit for gain control by behavioral state. Cell 156: 1139–1152. DOI: 10.1016/j.cell.2014.01.050, PMID: 24630718

Furutachi S, Franklin AD, Aldesa A, Mrsic-Flogel TD, Hofer SB. 2024. Cooperative thalamocortical circuit mechanism for sensory prediction errors. Nature 633, 398–406. DOi: 10.1038/s41586-024-07851-w. PMID: 39198646

Gămănuţ R, Kennedy H, Toroczkai Z, Ercsey-Ravaz M, Van Essen DC, Knoblauch K, Burkhalter A. 2018. The mouse cortical connectome, characterized by an ultra-dense cortical graph, maintains specificity by distinct connectivity profiles. Neuron 97: 698–715. DOI: 10.1016/j.neuron.2017.12.037, PMID : 29420935

Gao E, DeAngelis GC, Burkhalter A. 2010. Parallel input channels to mouse primary visual cortex. J Neurosci 30: 5912–5926. DOI: 10.1523/JNEUROSCI.6456-09.2010, PMID: 2042765

Goodale MA. 2011. Transforming vision into action. Vision research 51:1567–1587. DOI: 10.1016/j.visres.2010.07.027, PMID: 20691202

Gouwens NW, Sorensen SA, Berg J, Lee C, Jarsky T, Ting J, Sunkin SM, Feng D, Anastassiou CA, Barkan E, Bickley K, et al. 2019. Classification of electrophysiological and morphological neuron types in the mouse visual cortex. Nature Neurosci 22:1182–1195. DOI: 10.1038/s41593-019-0417-0, PMID: 31209381.

Guitchounts G, Masis J, Wolff SB, Cox D. 2020. Encoding of 3D Head orienting movements in primary visual cortex. Neuron 108: 512–525. DOI: 10.1101/2020.01.16.909473, PMID: 32783881

Han X, Bonin V. 2024. Higher-order cortical and thalamic pathways shape visual processing streams in ythe mouse cortex. Curr Biol 34: 1–14. DOI: 10.1016/j.cub.2024.10.048, PMID: 39566501

Harris JA, Mihalas S, Hirokawa KE, Whitesell JD, Choi H, Bernard A, Bohn P, Caldejon S, Casal J, Cho A, Feiner A, Feng D, Gaudreault N, Gerfen CR, Graddis N, Groblewski PA, Henry AM, Ho A, Howard R, Knox JE, Kuan X, Kuang X, Lecoq J, Lesnar O, Li Y, Luviano J, McConoughey S, Mortund MT, Naeemi M, Ng L, Oh SW, Ouellette B, Shen E, Sorensen SA, Wakeman W, Wang Q, Willford A, Phillips JW, Jones AR, Koch C, Zeng H. 2019. Hierarchical organization of cortical and thalamic connectivity. Nature 575: 195–202. DOI: 10.1038/s41586-019-1716-z, PMID: 31666704

Hilscher MM, Leão RN, Edwards SJ, Leão KE, Kullander K. 2017. Chrna2-Martinotti cells synchronize layer 5 types A pyramidal cells vis rebound excitation. PLOS Biology. DOI: 10.1371/journal.pbio.2001392, PMID: 28182735

Hoy JL, Niell CM. 2015. Layer-specific refinement of visual cortex function after eye opening in the awake mouse. Journal of Neuroscience 35: 3370–3383. DOI: 10.1523/JNEUROSCI.3174-14.2015, PMID: 25716837

Innocenti GM, Vercelli A. 2010. Dendritic bundles, minicolumns, columns, and cortical output units. Front Neuroanat 4: 11, DOI: https://doi.19.3389/neuro.05.011.2010, PMID : 20305751

Itokazu T, Hasegawa M, Kimura R, Osaki, H, Albrecht U-R, Sohya K, Chakrabarti S, Itoh H, Ito T, Sato TK, Sato TR. 2018. Streamlined sensory motor communication through cortical reciprocal connectivity in a visually guided eye movement task. Nat Commun 9: DOI: 10.1038/s41467-017-02501-4, PMID : 29362373

Jiang X, Wang G, Lee AJ, Stornetta RL, Zhu JJ. 2013. The organization of two new cortical interneuronal circuits. Nat Neurosci 16: 210–218. DOI: 10.1038/nn.3305, PMID: 23313910

Ji W, Gămănuţ R, Bista P, D’Souza RD, Wang Q, Burkhalter A. 2015. Modularity in the organization of mouse primary visual cortex. Neuron 87: 632–643. DOI: 10.1016/j.neuron.2015.07.004, PMID : 25247867

Jin M and Glickfeld LL. 2020. Mouse higher visual areas provide both distributed and specialized contributions to visually guided behaviors. Current Biology 30:4682–4692. DOI: 10.1016/j.cub.2020.09.015, PMID: 33035487

Juavinett AL, Kim EJ, Collins HC, Callaway EM. 2020. A systematic topographical relationship between mouse lateral posterior thalamic neurons and their visual cortical projection targets. J Comp Neurol 111: 528–599. DOI: 10.1002/cne.24737, PMID: 31265129

Kaneko M, Stryker MP. 2014. Sensory experience during locomotion promotes recovery of function in adult visual cortex. eLife 3: e02798. DOI: 10.7554/eLife.02798, PMID: 24970838

Karimi A, Odenthal J, Darwitsch F, Boergens KM, Helmstaedter M. 2020. Cell type specific innervation of cortical pyramidal cells at their apical dendrites. eLife 9: e46876. DOI: 10.7554/eLife.46876, PMID: 32108571

Keller AJ, Roth MM, Scanziani M. 2020a. Feedback generates a second receptive field in neurons of the visual cortex. Nature 582:545–594. DOI: 10.1038/s41586-020-2319-4. PMID: 32499655

Keller, AJ, Dipoppa M, Roth MM, Caudill MS, Ingrosso A, Miller KD, Scanziani M. 2020b. A disinhibitory circuit for contextual modulation in primary visual cortex. Neuron 108: 1181–1193.e8. DOI: 10.1016/j.neuron.2020.11.013, PMID : 33301712

Keller GB, Bonhoeffer T, Hübener M. 2012. Sensorimotor mismatch signals in primary visual cortex of the behaving mouse. Neuron 74: 809–815. DOI: 10.1016/j.neuron.2012.03.040, PMID : 22681686

Kohn A, Coen-Cagli R, Kanitscheider I, Pouget A. 2016. Correlations and neuronal population information. Annu Rev Neurosci 39: 237–256. DOI : 10.1146/annurev-neuro-07815-013851, PMID : 27145916

Kondo S, Yoshida T, Ohki K. 2016. Mixed functional microarchitectures for orientation selectivity in the mouse primary visual cortex. Nat. Commun. 7, 13210. DOI: 10.1038/ncomms13210. PMID: 27767032

Larkum ME, Waters J, Sakmann B, Helmchen F. 2007. Dendritic spikes in apical dendrites of neocortical layer 2/3 pyramidal neurons. J. Neurosci. 27: 8999–9008. DOI: 10.1523/JNEUROSCI.1717-07.2007, PMID: 17715337

Lee AM, Hoy JL, Bonci A, Wilbrecht L, Stryker MP, Niell CM. 2014. Identification of a brainstem circuit regulating visual cortical state in parallel with locomotion. Neuron 83: 455–466. DOI: 10.1016/j.neuron.2014.06.031, PMID : 25033185

Leinweber M, Ward DR, Sobczak JM, Attinger A, Keller GB. 2017. A sensorimotor circuit in mouse cortex for visual flow predictions. Neuron 95: 1420–1432. DOI: 10.1016/j.neuron.2017.08.036, PMID : 28910624

Mao D, Molina LA, Bonin V, McNaughton BL. 2020. Vision and locomotion combine to drive path integration sequences in mouse retrosplenial cortex. Curr. Biol. 30: 1680–1688.e4. DOI: 10.1016/j.cub2020.02.070, PMID: 32197086

Markov NT, Vezoli J, Chameau P, Falchier A, Quilodran R, Huissoud C, Lamy C, Misery P, Giroud P, Ullman S, Barone P, Dehay C, Knoblauch K, Kennedy H. 2014. Anatomy of hierarchy: feedforward and feedback pathways in macaque visual cortex. J Comp Neurol 522: 225–259. DOI: 10.1002/cne.23458, PMID: 23983048

Marshel JH, Garrett ME, Nauhaus I. Callaway EM. 2011. Functional specialization of seven mouse visual cortical areas. Neuron 72: 1040–1054. DOI: 10.1016/j.neuron.2011.12.004, PMID : 22196338

Maruoka H, Nakagawa N, Tsuruno S, Sakai S, Yoneda T, Hosoya T. 2017. Lattice system of functionally distinct cell types in the neocortex. Science 358: 610–615. DOI: 10.1126/science.aam6125, PMID : 29097542

Mazurek M, Kager M, Van Hooser SD. 2014. Robust quantification of orientation selectivity and direction selectivity. Front Neural Circuits 8: 92. DOI: 10.3389/fncir.2014.00092, PMID: 25147504

McBride EG, Lee S-YJ, Callaway EM. 2019. Local and global influences of visual spatial selection and locomotion in mouse primary visual cortex. Curr. Biol. 29: 1592–1605. DOI: 10.1016/jcub.2019.03.065, PMID: 31056388

Meier AM, Wang Q, Ji W, Ganachaud J, Burkhalter A. 2021. Modular network between postrhinal visual cortex, amygdala, and entorhinal cortex. J. Neurosci. 41: 4809–4825. DOI: 10.1523/JNEUROSCI.2185-20.2021, PMID: 33849948

Meyer HS, Egger R, Guest JM, Foerster R, Reissl S, Oberlaender M. 2013. Cellular organization of cortical barrel columns is whisker-specific. Proc Natl Acad Sci USA 110:19113–19118. DOI: 10.1073/pnas.1312691110, PMID: 24101458

Miura SK, Sanziani M. 2022. Distinguishing externally from saccade-induced motion in visual cortex. Nature 610: 135–142. DOI: 10.1038/s41586-022-05196-w. PMID: 36104560

Montijn JS, Vinck M, Pennartz CMA. 2014. Population coding in mouse visual cortex: response reliability and dissociability of stimulus tuning and noise correlation. Front Comput Neurosci. 8: 58. DOI: 10.3389/fncom.2014.00058. PMID: 24917812

Murayama M, Pérez-Garci E, Nevian T, Bock T, Senn W, Larkum ME. 2009. Dendritic encoding of sensory stimuli controlled by deep cortical interneurons. Nature 457:1137–1141. DOI: 10.1038/nature07663. PMID: 19151696

Musall S, Kaufman MT, Juavinett AL, Gluf S, Churchland AK. 2019. Single-trial dynamics are dominated by richly varied movements. Nat Neurosci 22:1677–1686. DOI: 10.1038/s41593-019-0502-4, PMID: 31551604

Niell CM, Stryker MP. 2010. Modulation of visual responses by behavioral state in mouse visual cortex. Neuron 65: 472–479. DOI: 10.1016/j.neuron.2010.01.033, PMID : 20188652

Pachitariu M, Stringer C, Dipoppa M, Schröder S, Rossi F, Dalgleish H, Carandini M, Harris KD. 2017. Suite2p: beyond 10,000 neurons with standard two-photon microscopy. bioRxiv.doi:10.1101/061507.

Pakan JM, Lowe SC, Dyla E, Keemink SW, Currie SP, Coutts CA, Rochefort NL. 2016. Behavioral-state modulation of inhibition is context-dependent and cell type specific in mouse visual cortex. eLife 5:e14985. DOI: 10.7554/eLife.14985. PMID: 27552056

Parker PRL, Abe ETT, Leonard ESP, Martins DM, Niell CM. 2022. Joint coding of visual input and eye/head position in V1 of freely moving mice. Neuron 110: 3897–3906. DOI: 10.1016/j.neuron.2022.08.029. PMID : 36137549

Paxinos G, Franklin KDJ. 2013. The Mouse Brain in Stereotaxic Coordinates. Forth Edition, Academic Press, Amsterdam.

Peters A, Kara DA. 1987. The neuronal composition of area 17 of rat visual cortex. IV. The organization of pyramidal cells. J Comp Neurol 260: 573–590. DOI: 10.1002/cne.902600410, PMID : 3611411

Polack PO, Friedman J, Golshani P. 2013. Cellular mechanisms of brain state-dependent gain modulation in visual cortex. Nat Neurosci 16:1331–1339. DOI: 10.1038/nn.3664, PMID : 23872595

Renart A, de la Rocha J, Bartho P, Hollender L, Parda N, Reyes A, Harris KD. 2010. The asynchronous state in cortical circuits. Science 327: 587–590. DOI: 10.1126/science.1179850. PMID: 20110507

Ringach DL, Mineault PJ, Tring E, Olivas ND, Garcia-Junco-Clemente P, Trachtenberg JT. 2016. Spatial clustering of tuning in mouse primary visual cortex. Nat Commun 7:12270. DOI: htps://doi.org/10.1038/ncomms12270. PMID: 27481398

Roth MM, Dahmen JC, Muir DR, Imhof F, Martini FJ, Hofer SB. 2016.Thalamic nuclei convey diverse contextual information to layer 1 of visual cortex. Nat. Neurosci. 19:299–301. DOI: 10.1038/nn.4197. PMID: 26691828

Schuman B, Dellal S, Prönneke A, Machold R, Rudy B. 2021. Neocortical layer 1: An elegant solution to top-down and bottom-up integration. Annu. Rev Neurosci. 44:221–252. DOI: 10.1146/annurev-neuro-100520-012117, PMID: 33730511

Shadlen NM, Newsome WT. 1998. The variable discharge of cortical neurons: implications for connectivity, computation, and information coding. J Neurosci 18: 3870–3896.DOI: 10.1523/NEUROSCI.18-10-03870.1998, PMID:9570816

Saleem AB, Ayaz A, Jeffery KJ, Harris KD, Carandini M. 2013. Integration of visual motion and locomotion in mouse visual cortex. Nat Neurosci 16: 1864–1869. DOI: 10.1038/nn.3567, PMID: 24185423

Salkoff DB, Zagha E, McCarthy E, McCormick DA. 2020. Movement and performance explain widespread cortical activity in a visual detection task. Cereb. Cortex 30: 421–437. DOI: 10.1093/cercor/bhz206, PMID:31711133

Sincich LC, Horton JC. 2002. Divided by cytochrome oxidase: a map of the projections from V1 to V2 in macaques. Science 295: 1734–1737. DOI: 10.1126.science.1067902. PMID: 11872845

Sincich LC, Horton JC. 2005. Input to V2 thin stripes arise from V1 cytochrome oxidase patches. J Neurosci 25:10087–10093. DOI: 10.1523/J.NEUROSCI.3313-05.2005, PMID: 16267215

Sit KK, Goard MJ. 2020. Distributed and retinotopically asymmetric processing of coherent motion in mouse visual cortex. Nat Commun 11:3565. DOI: 10.1038/s41467-020-17283-5. PMID: 32678087

Snyder LH, Grieve KL, Brotchie P, Andersen RA. 1998. Separate body-and world-references representations of visual space in parietal cortex. Nature 394, 887–890. DOI: 10.1038/29777, PMID: 9732870

Steinmetz NA, Zatka-Haas P, Carandini M, Harris KD. 2019. Distributed encoding of choice, action and engagement across the mouse brain. Nature 576: 266–273. DOI: 10.1038/s41686-019-1787-x, PMID: 31776518

Stringer C, Pachitariu M, Steinmetz N, Reddy CB, Carandini M, Harris KD. 2019. Spontaneous behaviors drive multidimensional, brainwide activity. Science 364: 255. DOI: 10.1126/science.aav7893, PMID : 31000656

Valente M, Pica G, Bondanelli G, Moroni M, Runyan CA, Morcos AS, Harvey CD, Panzeri S. 2021. Correlations enhance the behavioral readout of neural population activity in association cortex. Nat Neurosci 24: 975–986. DOI: https://10.1038/s441593-021-00845-1, PMID: 33986549

Vélez-Fort M, Bracey EF, Keshavarzi S, Rousseau CV, Cossell L, Lenzi SC, Strom M. and Margrie TW. 2018. A circuit for integration of head-and visual-motion signals in layer 6 of mouse primary visual cortex. Neuron 98:179–191. DOI: 10.1016/j.neuron.2018.02.023, PMID: 29551490

Vinck M, Batista-Brito R, Knoblich U. Cardin JA. 2015. Arousal and locomotion make distinct contributions to cortical activity patterns and visual encoding. Neuron 86:740–754. DOI: 10.1016/j.neuron.2015.03.028, PMID : 25892300

Wang A, Ferguson KA, Gupta J, Higley MJ, Cardin JA. (2024). Developmental trajectory of cortical somatostatin interneuron function. bioRxiv. 2024 Mar 7. DOI: 10.1101/2024.03.05.583530, PMID: 38496673

Wang Q, Sporns O, Burkhalter A. 2012. Network analysis of corticocortical connections reveals ventral and dorsal processing streams in mouse visual cortex. J Neurosci 32: 4386–4399. DOI: https://10.1523/JNEUROSCI.6063-11.2012, PMID: 22457489

Wekselblatt JB, Niell CM. 2019. Distinct functional classes of excitatory neurons in mouse V1 are differentially modulated by learning and task engagement. bioRxiv 533463. DOI: 10.1101/533463.

Williams SB, Arriaga M, Post WW, Korganokar AA, Moron JA, Han EB. 2019. Hippocampal activity dynamics during contextual reward association in virtual reality place conditioning. bioRxiv, DOI: 10.1101/54608

Wu SJ, Sevier E, Dwivedi D, Saldi G-A, Hairston A, Yu S, Abbott L, Choi DH, Sherer M, Qui Y, Shinde A, Lenahan M, Rizzo D, Xu Q, Barrera I, Kumar V, Marrero G, Prönneke A, Huang S, Kullander K, Stafford DA, Macosko E, Chen F, Rudy B, Fishell G. 2023. Cortical somatostatin interneuron subtypes for cell-type-specific circuits. Neuron 111: 2675–2692. DOI: 10.1016/j.neuron.2023.05.032. PMID : 37390821

Yang Y, Lee J, Kim G. 2020. Integration of locomotion and auditory signals in the mouse inferior colliculus. eLife 9: e52228. DOI: 10.7554/eLife.52228. PMID: 31987070

Yu Y, Stirman JN, Dorsett CR, Smith SL. 2019. Mesoscale correlation structure with single cell resolution during visual coding. Preprint at 10.1101/469114

Yu Y, Stirman JN, Dorsett CR, Smith SL. 2022. Selective representations of texture and motion in mouse visual areas. Curr Biology 32:2810–2820. DOI: 10.1016/j.cub.2022.04.091, PMID: 35609609

Zhang S, Xu M, Kamigaki T, Do JPH, Chang W-C, Jenvay S, Miyamichi K,Luo L, Dan Y. 2014. Long-range and local circuits for top-down modulation of visual cortex processing. Science 345:660–665. DOI: 10.1126/science.1254126. PMID: 25104383

